# Multi-omic profiling of human and mouse dorsal root ganglia enables targeted gene delivery to nociceptors

**DOI:** 10.64898/2026.03.05.709931

**Authors:** Lily S. He, Parth Bhatia, Shamsuddin A. Bhuiyan, Evangelia Semizoglou, Jiaxiang Wang, Jia Li, Joohyun Nam, Jay X. J. Luo, Chayse Arnhold, Difei Zhu, Mengyi Xu, Dustin Griesemer, Hyo Jeong Yong, Lorna Jayne, Evangeline Gilmer, Qiyi Li, Katerina Pantaleo, Lite Yang, Erika K. Williams, Selwyn Jayakar, Brian J. Wainger, Sinisa Hrvatin, William Renthal, NIH PRECISION Human Pain Network

## Abstract

Chronic pain conditions are often driven by hyperexcitability of nociceptors, the peripheral sensory neurons that detect noxious stimuli. Recent single-cell transcriptomic studies have begun to clarify the molecular identity of distinct peripheral sensory neuron subtypes, but tools that restrict transgene expression to nociceptors while sparing other dorsal root ganglion (DRG) subtypes remain limited. Here, we combined single-nucleus multi-omic profiling of human and mouse DRG with *in vivo* AAV enhancer screening to identify *cis*-regulatory elements that drive biased AAV expression in mouse DRG nociceptors and human iPSC-derived nociceptors. We then validated that an enhancer AAV designed to express K_ir_2.1 preferentially in nociceptors reduces DRG neuronal excitability. Leveraging these multi-omic datasets, we trained a sequence-based model to decode the *cis*-regulatory logic of nociceptors, enabling both the prioritization of native candidate elements and the design of synthetic enhancers with a range of nociceptor targeting properties. Together, these cross-species multi-omic resources define conserved DRG regulatory programs and provide a viral toolkit for pain research with potential translational applications for patients with refractory pain.

## Introduction

Chronic pain is often driven by maladaptive hyperexcitability of nociceptors, which are a class of peripheral sensory neurons that detect noxious stimuli. Most analgesics act on multiple cell types across central and peripheral nervous systems, which in the case of opioids, can lead to potentially dangerous side effects.^1–4^ Thus, there is an urgent need for more targeted, cell-type-specific strategies to modulate nociceptor activity selectively while sparing other neuron types. Toward that goal, recent single-cell/nucleus RNA sequencing (sc/snRNA-seq) studies of DRG have revealed unique transcriptomic profiles of multiple nociceptor subtypes that are largely conserved between rodents and humans.^5,6^ Transgenic mice engineered to target distinct DRG subtypes have yielded important insights into their function,^7–13^ but these methods are not easily translated to other model organisms or humans. Indeed, there are currently no tools that provide subtype-selective access to nociceptors across species.

Recent advances in single-cell multi-omic technologies offer a new strategy for characterizing the gene-regulatory logic underlying DRG cell-type-specific gene expression. By jointly profiling chromatin accessibility and gene expression in the same nucleus, it is possible to identify *cis*-regulatory elements (CREs) that promote cell-type-specific gene expression within defined neuronal populations. Engineering AAVs to use cell-type-specific CREs for driving transgene expression in defined cell populations has been successfully implemented in the brain and spinal cord,^14–17^ but it has been challenging to apply this approach to peripheral sensory neurons due to our limited understanding of the DRG neuronal epigenome.

Here, we constructed DRG multi-omic cell atlases in both human and mouse and utilized joint chromatin accessibility and transcriptomic data to define evolutionarily conserved *cis*-regulatory programs and CREs that are preferentially active in nociceptors. We built a suite of enhancer

AAVs and quantified their ability to drive viral expression preferentially in nociceptors *in vivo* using scRNA-seq and spatial transcriptomics. We then trained a sequence-based model to decode the *cis*-regulatory logic of nociceptors, which can be used to prioritize native CREs for enhancer AAVs and to design synthetic enhancers. Finally, we show that an enhancer AAV that preferentially expresses in mouse nociceptors relative to other DRG cell types significantly reduces DRG neuronal excitability in mice and drives strong expression in human iPSC-derived nociceptors. Together, these findings establish a cross-species DRG multi-omic resource and outline a general strategy for developing cross-species viral tools for translational pain research with potential therapeutic applications.

## Results

### Single-nucleus multi-omic profiling of mouse and human DRG

To characterize the *cis*-regulatory logic of mouse and human DRG cell types, we constructed multi-omic (snRNA-seq and snATAC-seq) cell atlases of DRG from 71 male and female adult mice and 66 human donors (32 male, 34 female; Table S1) (Figures 1A). This multi-omic approach allows for the simultaneous interrogation of transcriptional output and chromatin accessibility within the same nucleus, enabling the direct linkage of distal CREs to their target genes without the ambiguity of *in silico* integration.^18^ As DRG neurons comprise only ∼1% of total cells in the ganglion,^5,6^ we optimized neuronal enrichment protocols for both mouse and human DRG prior to multi-omic profiling. In mice, we sorted genetically labeled DRG neurons (*Vglut2*-Cre+/-; *Sun1*-GFP+/-) and observed that >90% of DRG cells sequenced are neurons (Figure S1A). In humans, we sorted NeuN-stained nuclei and observed that up to 60% of DRG cells sequenced are neurons (Figure S1B). We then performed snMultiome of neuronally enriched mouse and human DRG samples, and after quality controls (see Methods, Figure S1C-D, S2A-D), we obtained 23,928 mouse DRG nuclei, including 12,272 epigenomically profiled nuclei from prior work,^19,20^ and 155,008 human DRG nuclei. We next performed unsupervised clustering based on gene expression and annotated cell types using the expression of known marker genes (see Methods).^5,6^ We annotated eight broad cell types in both mouse and human DRG: neurons, fibroblasts, satellite glia (SGC), myelinating Schwann Cells (mySC), non-myelinating Schwann Cells (nmSC), endothelial cells, immune cells, and pericytes (Figures 1B-C; Figure S2E-G). Gene expression and chromatin accessibility at loci of known marker genes were consistent with anchored cell type annotations (Figures 1B-C, bottom).^5,6^ and largely similar between donors and sexes (Figures S3A-D). Differential gene expression between DRG cell types identified 9,856 differentially expressed genes in mice and 11,777 in humans (log_2_FC > 0.5, adj. p-val < 0.05, cell type compared to all others) (Table S2 and S3, see Bhuiyan et al., co-submitted for in depth gene expression analyses of human DRG cell types).

**Figure 1:**
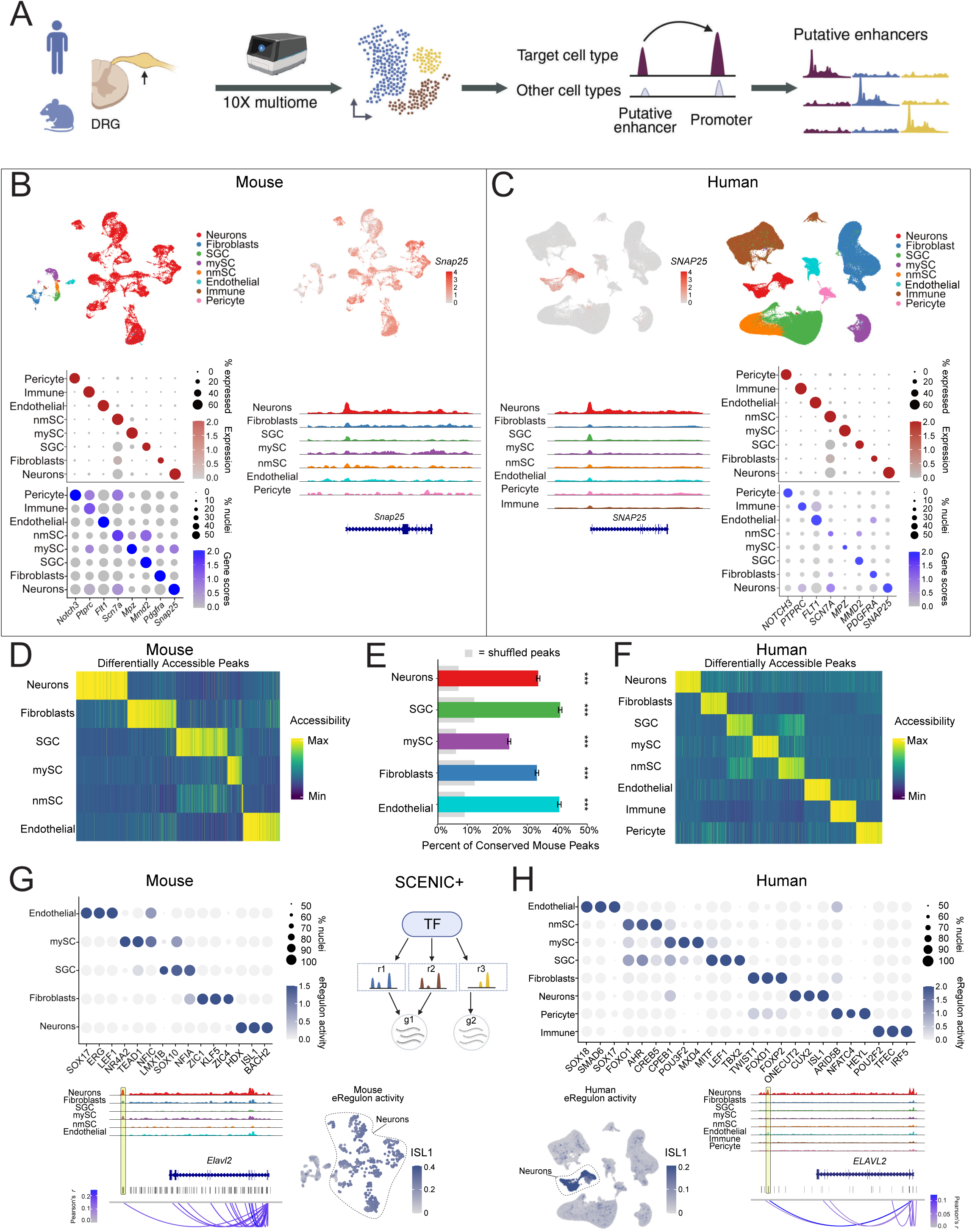
snMultiome of mouse and human DRG. **A:** Workflow displays schematic of experimental design. **B:** (*Top, Left*) UMAP projections displaying 23,928 mouse DRG nuclei colored by cell type or (*Top, Right*) log-normalized *Snap25* expression. (*Bottom, Left*) Dot plot displaying log-normalized gene expression (red) or log-normalized ATAC gene activity scores (blue) for DRG cell type marker genes. ATAC gene activity scores are defined as the sum of ATAC fragments falling within the gene body and the 2 kb region upstream of the transcription start site (TSS). Dot size represents the percentage of nuclei expressing the marker gene (RNA) or the percentage of nuclei with accessible chromatin at the marker gene locus (ATAC). (*Bottom, Right*) Genomic tracks displaying normalized chromatin accessibility at the *Snap25* locus for each mouse DRG cell type. **C:** (*Top, Left*) UMAP projections displaying 155,008 human nuclei colored by log-normalized *SNAP25* expression or (*Top, Right*) cell type. (*Bottom, Left)* Genomic tracks displaying normalized chromatin accessibility at the *SNAP25* locus for each human DRG cell type. (*Bottom, Right*) Dot plot displaying log-normalized gene expression (red) or log-normalized ATAC gene activity scores (blue) for canonical marker genes. ATAC gene activity scores are defined as the sum of ATAC fragments falling within the gene body and the 2 kb region upstream of the TSS. Dot size represents the percentage of nuclei expressing the marker gene (RNA) or the percentage of nuclei with accessible chromatin at the marker gene locus (ATAC). **D:** Heatmap displays the average (Z-scaled Transverse Frequency-Inverse Document Frequency [TF-IDF] normalized mean) number of fragments for all nuclei across the top 2,000 differentially accessible peaks (columns) for each mouse DRG cell type (log_2_FC > 0.5, adj. p-val < 0.05 relative to all other cell types). Rows represent DRG cell types. **E:** Bar chart displays the mean proportion of mouse differentially accessible (DA) peaks (log_2_FC > 0.5, adj. p-val < 0.05 relative to all other mouse cell types) that are syntenic with human DRG differentially accessible peaks (log_2_FC > 0.5, adj. p-val < 0.05, relative to all other human cell types) in the corresponding DRG cell type. Values are estimated by bootstrapping (100 iterations), sampling 50 peaks per iteration from the top 500 mouse DA peaks for each cell type. Error bars indicate standard error across iterations. Gray bars indicate control test done using differentially accessible peaks with randomly shuffled cell type labels (*** = adj.p-value < 0.0001, Z-test comparing distribution of overlap of differentially accessible peaks with peaks randomly shuffled across cell types). **F:** Heatmap displays the average (Z-scaled TF-IDF-normalized mean) number of fragments for all nuclei across the top 2,000 differentially accessible peaks (columns) for each human DRG cell type (log_2_FC > 0.5, adj. p-val < 0.05 relative to all other cell types). Rows represent DRG cell types. **G:** (*Top, Left*) Dotplot displays the eRegulon activity scores (dot color) of transcription factors in mouse DRG cell types. Dot size represents the percentage of nuclei with non-zero eRegulon scores. (*Top, Right*) Diagram displays schematic of what is factored in the calculation of an eRegulon activity score - transcription factor (TF) expression, peak region accessibility and target gene expression. (*Bottom, Left*) Genomic tracks display normalized chromatin accessibility at the *Elavl2* locus for each mouse DRG cell type. Highlighted region indicates a candidate *cis*-regulatory element (CRE) with a predicted ISL1 binding motif. Loops show Pearson correlation (p-val. < 0.05) of CRE accessibility and RNA expression of *Elavl2.* (*Bottom, Right*) UMAP projection displays ISL1 eRegulon activity scores (color scale) in mouse DRG cell types (downsampled to 2,414 nuclei for visualization purposes). **H:** (*Top*) Dotplot displays the eRegulon activity scores (dot color) of transcription factors expressed in human DRG cell types. Dot size represents the percentage of nuclei with non-zero eRegulon scores. (*Bottom, Left*) UMAP projection displays ISL1 eRegulon activity scores (color scale) across 95,515 human nuclei and includes all cell types found in Fig. 1C. (*Bottom, Right*) Genomic tracks display normalized chromatin accessibility at the *ELAVL2* locus for each human DRG cell type. The highlighted region indicates a candidate CRE with a predicted ISL1 binding motif. Loops show Pearson correlation (p-val. < 0.05) of CRE accessibility and RNA expression of *ELAVL2*.

To identify regions of transposase-accessible chromatin in each DRG cell type, we performed “peak” calling (see Methods) and observed that peaks of accessible chromatin are predominantly located in intronic and non-coding regions distal to the gene body in both mouse and human (Figures S3E-F; see Methods). We then performed differential chromatin accessibility analysis for the 8 broad DRG cell types by comparing peaks of accessible chromatin in one cell type to all other cell types. We observed 268,405 unique differentially accessible peaks in mice and 252,806 in humans (log_2_FC > 0.5, adj. p-val < 0.05, cell type compared to all others) (Figure 1D-F; Table S4 and S5). To identify overlap in differentially accessible peaks between the two species, we used the LiftOver tool to identify human regions with high sequence similarity (syntenic) to the mouse differentially accessible peaks.^21,22^ Among these syntenic regions, we observed significant overlap (35-44%) between the differentially accessible peaks in corresponding mouse and human DRG cell types relative to a null distribution generated by shuffling cell type identities within the pool of top accessible regions (adj. p-value <10^-80^, Fisher’s Exact test) (Figure 1E). This degree of overlap is consistent with conservation of cell-type-specific regulatory programs between mouse and human DRG, and suggests that some differentially accessible peaks may mark shared CREs across species.^23^

We next used SCENIC+ to characterize the gene regulatory networks of transcription factors (TFs), CREs, and predicted target genes that establish cell identity in mouse and human DRG cell types. SCENIC+ quantifies the activity of “eRegulons” in each cell by integrating co-expression of a given transcription factor (TF) and predicted target genes with chromatin accessibility and TF motif enrichment at corresponding putative CREs.^24^ We observed multiple TFs in which their eRegulons are preferentially active in distinct DRG cell types compared to all other DRG cell types in both mouse (Figure 1G; Table S6) and human (Figure 1H; Table S7). For example, *ISL1*, a transcription factor known to regulate DRG neuronal development,^25^ displays significantly higher eRegulon activity in DRG neurons than in non-neuronal cell types in both mouse and human. This ISL1 eRegulon includes the mature neuronal marker gene, *Elavl2*, and a syntenic CRE at which there is both an ISL1 consensus binding site and increased chromatin accessibility in both mouse and human neurons compared to other DRG cell types (Figures 1G-H). Consistent with observations in central nervous system cell types,^15,26–30^ the conserved, cell-type-specific regulatory activity of certain TFs between mouse and human DRG cell types suggests that CREs identified in mice have the potential to recruit similar TFs and confer comparable regulatory activity in humans.

### Cross-species cis-regulatory logic of nociceptors

To identify regulatory elements that are selectively active in nociceptors across species, we next subclustered 21,064 mouse and 6,893 human DRG neurons by their gene expression profiles (Tables S8-9) and annotated their neuronal subtype by anchoring to previous human and mouse DRG sc/snRNA-seq studies.^6,31–33^ We observed the same ∼18-22 DRG neuronal subtypes in both mouse and human that have been reported previously (Figure 2A), which includes the important C-peptidergic (C-PEP) and C-non-peptidergic (C-NP) nociceptors that are implicated in chronic pain conditions.^6,31–33^ After filtering the annotated mouse and human neuronal nuclei for those with high quality chromatin accessibility data (Figure S1C-D; Figure S4A-B; see Methods), we performed downstream epigenomic analysis on 18,238 mouse and 1,597 human DRG neurons. As human DRG neurons are more challenging to obtain high quality chromatin accessibility data than in mouse or even in DRG non-neuronal cells (Figure S4C-D), we grouped human neuronal subtypes into 6 broader classes (A-Propr, A-LTMR, A-PEP, C-PEP, C-NP, and C-Thermo) for downstream epigenomic analyses. We observed neuronal subtype-specific patterns of chromatin accessibility at corresponding marker gene loci for each of these neuronal classes (Figure 2B-E; Figure S4E-F). Utilizing all DRG neurons that passed quality controls in mouse or human (Figure S4C-D), we next performed genome-wide differential chromatin accessibility analysis across distinct DRG neuronal subtypes in each species. We identified 575,462 peaks in mice and 9,370 peaks in humans that are differentially accessible (log_2_FC > 0.5, adj. p-val < 0.05 [mouse], < 0.1 [human]) in one DRG neuronal subtype compared to the others (Figures 2C, 2E, Tables S10 and S11). The difference in numbers of differentially accessible peaks between mouse and human is largely driven by their respective neuron numbers; downsampling mouse DRG neuron numbers to match human neuron numbers yielded ∼12,000 differentially accessible peaks (log_2_FC > 0.5, adj. p-val < 0.05). Differentially accessible peaks across mouse DRG neuronal subtypes are significantly more likely to overlap with differentially accessible peaks across human DRG neuronal subtypes compared to random length-matched ENCODE CREs (Fisher’s exact test, p < 10^-16^), supporting some conservation of chromatin accessibility between mouse and human DRG neurons.^34^

**Figure 2:**
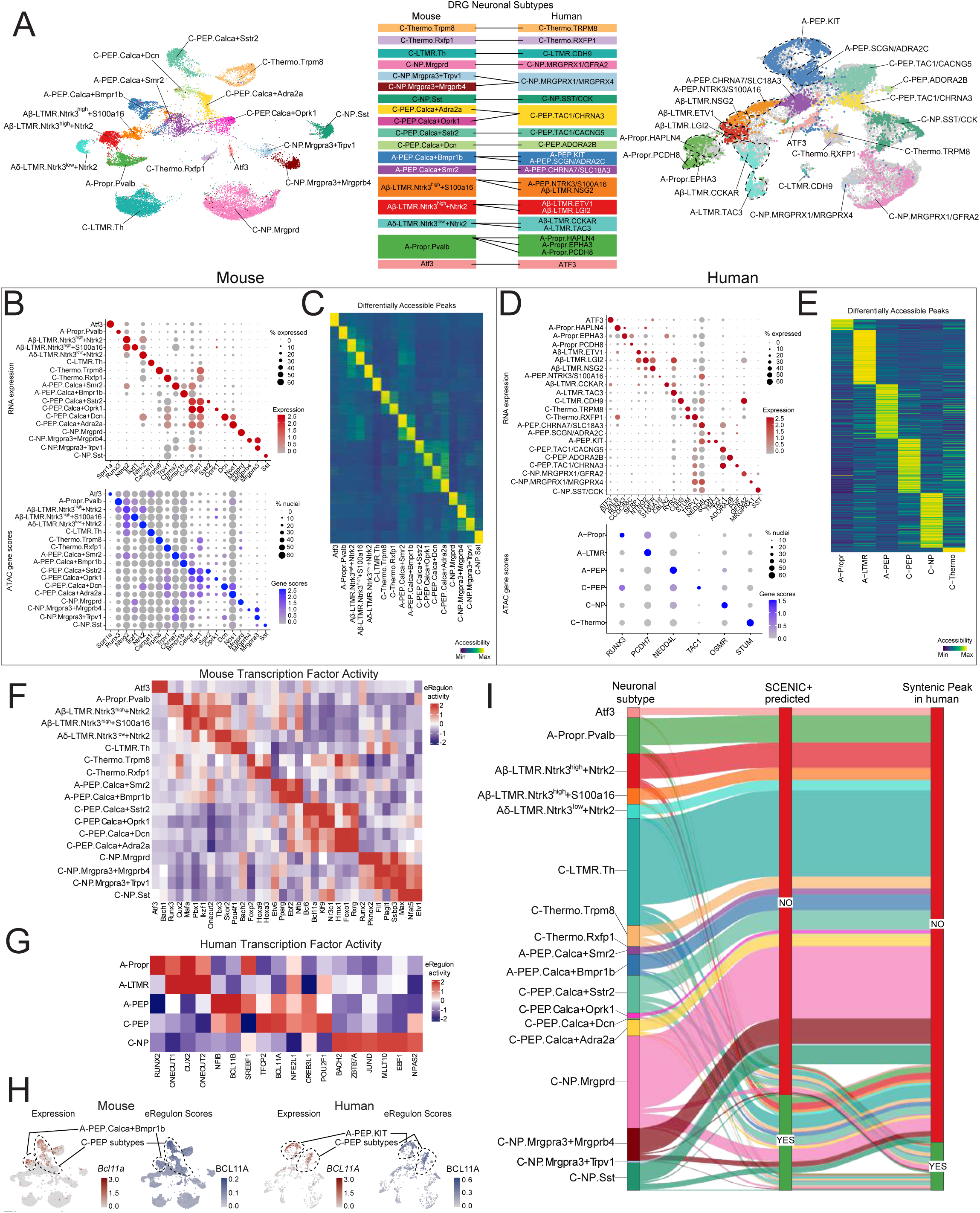
snMultiome of mouse and human DRG neuronal subtypes. **A:** UMAP projection displays snMultiome data from 21,064 mouse (*Left*) and 6,893 human (*Right*) DRG neurons. Nuclei are projected into the mouse^6^ or human^5^ reference atlas space. (*Middle*) Nuclei are colored by neuronal subtype, according to the key between the UMAPs. Transcriptomically corresponding neuronal subtypes between mouse and human are connected by lines. **B:** Dot plot displaying log-normalized gene expression (red) or log-normalized ATAC gene activity scores (blue) for marker genes of mouse DRG neuronal subtypes. ATAC gene activity scores are defined as the sum of ATAC fragments falling within the gene body and the 2 kb region upstream of the TSS. Dot size represents the percentage of nuclei expressing the marker gene (RNA) or the percentage of nuclei with accessible chromatin at the marker gene locus (ATAC). **C:** Heatmap displays the average (Z-scaled TF-IDF-normalized mean) number of fragments for all nuclei across the top 2,000 differentially accessible peaks (rows) for each mouse neuronal subtype (log_2_FC > 0.5, adj. p-val < 0.05 relative to all other neuronal subtypes). Columns represent DRG neuronal subtypes. **D:** Dot plot displaying log-normalized gene expression (red) or log-normalized ATAC gene activity scores (blue) for canonical marker genes for human neuronal subtypes. ATAC gene activity scores are defined as the sum of ATAC fragments falling within the gene body and the 2 kb region upstream of the TSS. Dot size represents the percentage of nuclei expressing the marker gene (RNA) or the percentage of nuclei with accessible chromatin at the marker gene locus (ATAC). **E:** Heatmap displays the average (Z-scaled TF-IDF-normalized mean) number of fragments for all nuclei across the top 150 differentially accessible peaks (rows) for each human neuronal subtype (log_2_FC > 0.5, adj. p-val < 0.1 relative to all other neuronal subtypes). Columns represent DRG neuronal subtypes. **F-G:** Heatmap displays SCENIC+ eRegulon activity scores (z-scaled) of TFs (columns) in mouse (F) or human (G), averaged across DRG neuron subtypes (rows). **H:** UMAP projections display log-normalized *Bcl11a* gene expression (first UMAP from the left) and BCL11A eRegulon activity scores (second UMAP) in mouse DRG neuronal nuclei. UMAP projections displaying log-normalized *BCL11A* expression (third UMAP) and BCL11A SCENIC+ eRegulon activity scores (fourth UMAP) for human DRG neuronal nuclei are also shown. **I:** Sankey plot displays differentially accessible peaks in mouse for each neuronal subtype (log_2_FC > 0.5, adj. p-value < 0.05 relative to all other neuronal subtypes) that are also predicted to comprise SCENIC+ eRegulons or which have syntenic peaks in the human DRG neurons. Bars and ribbons are colored by mouse neuronal subtypes. Green vertical bars indicate overlap, red vertical bars indicate no overlap.

We next used SCENIC+ to prioritize differentially accessible peaks that may function as neuronal subtype-specific CREs in both humans and mice. In mice, SCENIC+ nominated 484 ‘eRegulons’ and 61,044 putative CREs, and in humans, SCENIC+ nominated 53 eRegulons and 3,384 putative putative CREs (Figures 2F-G, Tables S12 and S13). Clustering DRG neurons by their eRegulon activities was sufficient to resolve known neuronal identities (Figure S4G), highlighting the potential of these inferred gene regulatory networks to identify key TFs and candidate CREs that drive neuronal subtype-specific gene expression in both mouse and human. For example, SCENIC+ analyses identified increased RUNX3 eRegulon activity in proprioceptors compared to other neuronal subtypes and a putative CRE upstream of the proprioceptor-specific gene, *Pvalb*, which is consistent with the known role of RUNX3 in proprioceptor development (Figures 2F, S4H).^35,36^ In C-PEP nociceptors, SCENIC+ implicates BCL11A with increased eRegulon activity compared to other neuronal subtypes in both mouse and human and nominates a putative CRE upstream of the nociceptor-enriched gene, *Sstr2* (Figures 2F-G, S4H).

These findings suggest that our mouse and human DRG multi-omic atlases and downstream gene regulatory network analyses could help identify CREs that are capable of driving nociceptor-specific gene expression in both species, a critical requirement for developing translational viral tools with targeted activity in nociceptors. Toward that goal, we found that ∼16% of all differentially accessible peaks between mouse neuronal subtypes are also linked to a SCENIC+ eRegulon and that ∼18% of these have syntenic peaks in human DRG neurons (Figure 2I; Table S14). This analysis highlights a core set of CREs that are implicated in cell-type-specific gene regulation and have the potential to function similarly across species.

### In vivo screening of candidate nociceptor-specific CREs

To functionally screen for CREs that drive gene expression preferentially in nociceptors *in vivo*, we constructed AAVs in which prioritized CREs are inserted upstream of a minimal promoter (mini CMV) and a reporter gene (EGFP). AAVs were then delivered to mice intracerebroventricularly (ICV), and reporter expression was quantified in each DRG neuronal subtype using fluorescent *in situ* hybridization (FISH) and/or scRNA-seq (Figure 3A). We initially selected 9 CREs (Figure 3B; see Methods) based on their differential accessibility patterns in two important classes of nociceptors in pain transduction, C-PEP (CRE1-7) or C-NP (CRE8-9) nociceptors. We also included two controls: an AAV with only a minimal promoter driving reporter expression (No CRE) and an AAV with the ubiquitous CAG promoter driving reporter expression. The 11 AAVs (9 nociceptor CREs, 2 controls) were packaged in the AAV-PHP.S capsid^37^ and injected ICV into neonatal mice to ensure broad DRG infectivity;^38–40^ DRGs were collected at least 6 weeks post-injection.

**Figure 3:**
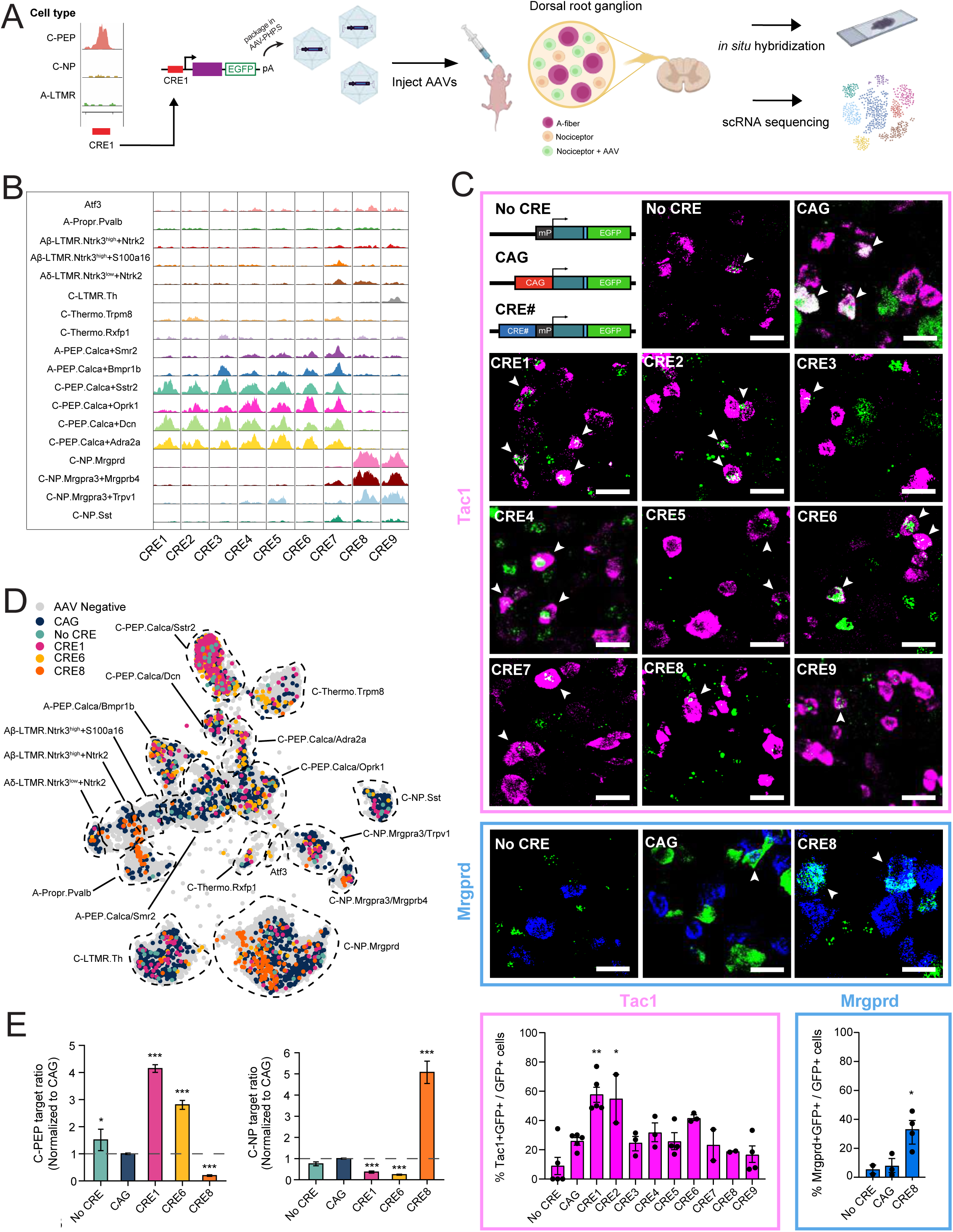
Screening candidate CREs for nociceptor specificity. **A:** Schematic of the experimental strategy used to screen AAV expression patterns in DRG driven by candidate CREs. **B:** Genomic tracks display chromatin accessibility at the loci of candidate CREs in distinct mouse DRG neuronal subtypes. **C:** (*Top, Left*) Viral cargo schematic (mP: minimal promoter, teal box: reporter construct, blue box: P2A) and representative 20X images of mouse DRG labeled by RNAscope *in situ* hybridization for *Tac1* (magenta) and *EGFP* (green) mRNA (first 4 rows), or for *Mrgprd* (blue) and *EGFP* (green) mRNA (bottom row). White arrowheads indicate either *Tac1* and *EGFP* or *Mrgprd* and *EGFP* signal colocalization. (*Bottom, Left*) Quantification of *Tac1* and *EGFP* mRNA colocalization for each candidate CRE. (One-way ANOVA, F(3,27) = 6.80, p < 0.001; Dunnett’s multiple comparisons test; **p = 0.001, CRE1 vs. CAG, * p < 0.05, CRE2 vs. CAG). (*Bottom, Right*) Quantification of *Mrgprd* and *EGFP* mRNA colocalization for CRE8 (One-way ANOVA, F(2,6) = 7.42, p = 0.024; Dunnett’s multiple comparisons test; *p < 0.05 CRE8 vs. CAG). Data are presented as mean ± SEM. Each point represents a biological replicate. n = 2-5 animals per CRE). Scale bar, 25 µm. **D:** UMAP projection displays 34,899 mouse DRG neurons, 1,032 of which expressed AAV transcripts (>1 AAV transcript/cell; colored by CREs). Cells without detected AAV transcripts are gray. Projections and cell type annotations use the mouse reference UMAP space.^6^ **E:** Quantification of C-PEP (*Left*) or C-NP (*Right*) target ratios (AAV transcripts in C-PEP or C-NP subtype / AAV transcripts in all other subtypes) for candidate CRE1, CRE6, and CRE8 as measured by scRNA-seq. Cell class specificity target ratios are normalized to the CAG control ratio. Error bars indicate the standard deviation of randomly sampled cells in each group (n = 5 iterations). Significance was determined by Fisher’s exact test (ns: not significant, *: p < 0.05, **: p < 0.01, ***: p < 0.001). C-PEP subtypes: C-PEP.Calca+Sstr2, C-PEP.Calca+Dcn, C-PEP.Calca+Oprk1; C-NP subtypes: C-NP.Mrgprd, C-NP.Mrgpra3+Trpv1, C-NP.Mrgpra3+Mrgprb4.

We used FISH to quantify co-expression of viral eGFP with either the C-PEP nociceptor marker, *Tac1*, or the C-NP nociceptor marker, *Mrgprd*. Two of the nine tested C-PEP nociceptor CREs (CRE1, CRE6) drove reporter expression with significantly greater specificity in *Tac1*+ neurons than the CAG control (Figure 3C) (One-way ANOVA, F(3,27) = 6.80, p < 0.001; Dunnett’s multiple comparisons test). We additionally confirmed that CRE1, which displayed the greatest specificity for C-PEP by FISH, drives limited expression in off-target cell types (*Mrgprd*+ C-NP nociceptors, *Hapln4*+ Aβ-LTMRs and proprioceptors, and *Fam19a4*+ C-LTMRs) (Figure S5A). We also identified one of the C-NP targeting CREs, CRE8, drives eGFP expression significantly more specifically in *Mrgprd*+ C-NP neurons than the CAG control (Figure 3C) (One-way ANOVA, F(2,6) = 7.42, p = 0.024; Dunnett’s multiple comparisons test).

As neither *Tac1* nor *Mrgprd* marker genes label all C-PEP or C-NP nociceptors, respectively (Figure 3, S5A), we turned to scRNA-seq to provide an orthogonal and more quantitative assessment of AAV reporter expression for CREs that displayed promising nociceptor specificity by FISH. In particular, CRE1, CRE6, CRE8, and control (No CRE, CAG) AAVs were delivered ICV to neonates as described above, followed by whole cell dissociation of DRGs and scRNA-seq at least 6 weeks post-injection. In total, 34,899 cells were profiled by scRNA-seq across all CREs and controls, with 2-4 biological replicates per CRE. Neuronal subtype annotations were obtained by anchoring scRNA-seq data to the mouse DRG reference atlas^6^ (Figure 3D) and AAV transcripts were quantified in each neuronal subtype. C-PEP target specificity ratios (AAV+ C-PEP cells / all other AAV+ cells) and C-NP target specificity ratios (AAV+ C-NP cells / all other AAV+ cells) were calculated for each CRE and compared to control AAVs. Consistent with the FISH results, we observed that CRE1 and CRE6 drive reporter expression more specifically in C-PEP nociceptors than No CRE or CAG controls (∼3-5 fold, Fisher’s exact test: p < 0.001), and CRE8 drives reporter expression more specifically in C-NP nociceptors than No CRE or CAG controls (∼4-6 fold, Fisher’s exact test: p < 0.001) (Figure 3E). The additional granularity of scRNA-seq revealed that, of C-PEP nociceptors (C-PEP.Calca+Sstr2, C-PEP.Calca+Oprk1, C-PEP.Calca+Dcn), both CRE1 and CRE6 drive expression preferentially in C-PEP.Calca+Sstr2 neurons compared to other C-PEP subtypes and that, of C-NP nociceptors (C-NP.Mrgprd, C-NP.Mrgpra3+Trpv1, C-NP.Mrgpra3+Mrgprb4), CRE8 drives reporter expression preferentially in C-NP.Mrgprd neurons compared to other C-NP subtypes (Figure S5B). These scRNA-seq results were orthogonally assessed by FISH (Figure S5C), where we observed that 56 ± 2% of CRE1 and 45 ± 13% of CRE6 driven reporter positive cells were C-PEP.Calca+Sstr2 compared to 28 ± 3% of No CRE and 36 ± 1% of CAG controls, respectively (Figure S5D) (One-way ANOVA, F(3,8) = 2.005, p = 0.0404, Dunnett’s multiple comparisons test for No CRE versus CRE1. Data are presented as mean ± SEM).

Given the low success rate and high cost of screening individual CREs, we turned to a multiplexed AAV screen to functionally assess CRE activity *in vivo* more efficiently (Figure 4A).^41^ We selected 106 CREs predicted to drive expression in C-PEP nociceptors, C-NP nociceptors, *Sst+* pruriceptors, or *Mrgpra3+* pruriceptors, or combinations of nociceptor classes, and constructed an AAV library in which each CRE was cloned in *cis* with unique DNA barcodes (see Methods). We also included a No CRE control that was uniquely barcoded. Next-generation sequencing of the resulting AAV library confirmed that all CREs were represented (Figure S6A). After packaging the CRE library in AAV-PHP.S, the AAV library was injected ICV into three neonatal mice and scRNA-seq of their DRGs was performed >6 weeks later as described above. After quality control, clustering, and cell type annotation, a total of 10,442 DRG cells were included for downstream analysis. Of these, 2,275 cells expressed AAV transcripts covering 106 CREs (Figure 4B), of which 41 CREs were detected in at least 6 cells and included in downstream analyses (Figure S6B). We observed that certain CREs drove barcode expression preferentially in distinct DRG neuronal cell classes compared to the No CRE control (Figure 4C), with scRNA-seq providing the resolution to identify expression within specific subtypes (Figure S6C). For example, CREs accessible in peptidergic neurons (Figure 4C) drove expression in C-PEP (CRE70 and CRE71) or A-PEP (CRE18) nociceptors (Figure S6C). Similarly, CREs accessible in non-peptidergic cells drove expression in C-NP.Mrgpra3 pruriceptors (CRE45 and CRE47) or C-NP.Sst pruriceptors (CRE114). This specificity extended to non-nociceptive lineages, where CRE68 and CRE79 drove preferential expression in A-Propr.Pvalb and C-LTMR.Th cell types, respectively.

**Figure 4:**
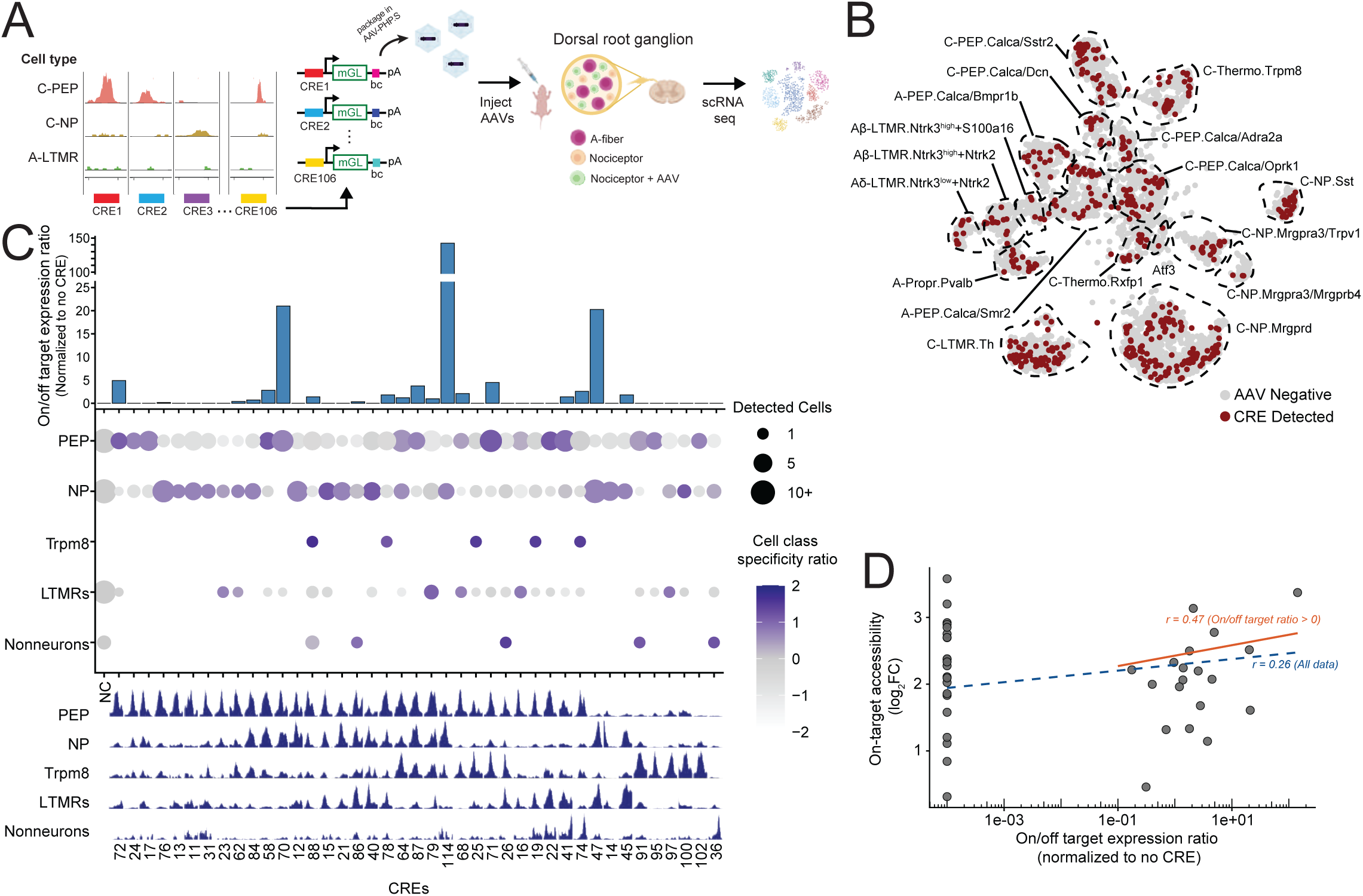
Multiplexed **in vivo** AAV screen to functionally assess CRE activity in DRG. **A:** Schematic of the experimental strategy used for multiplexed CRE screen. mGL: mGreenLantern; bc: Barcode. **B:** UMAP projection displays scRNA-seq of 10,442 DRG cells from mice infected with a multiplexed AAV CRE library. Cells are colored if AAV barcodes were detected (>1 barcode/cell; 2,275 cells). Projections are in the mouse reference UMAP space.^6^ **C:** (*Bottom*) Genomic tracks display chromatin accessibility profiles in mouse DRG at loci for candidate CREs included in specificity analysis across 5 DRG cell classes. (*Middle*) Dotplot displays cell class specificity ratios across 5 DRG cell classes. The cell class specificity ratio (y-axis) is defined as the number of CRE barcode-positive cells in a specific cell class divided by the number of barcode-positive cells in all other cells. Dot color represents cell class specificity ratio (Z-scaled), and dot size corresponds to the number of a given CRE’s barcodes that were detected per cell type. Ratios are normalized to the negative control (NC: No CRE). (*Top*) The bar plot displays the on/off target expression ratio for each CRE, where on-target cell types are defined by the chromatin accessibility at each CRE locus. The on/off target expression ratio is calculated as the number of CRE barcodes detected in on-target cell types cell types divided by the number of CRE barcodes cells in off-target (inaccessible) cell types, normalized to the negative control (No CRE). Only CREs with viral barcodes detected in > 6 cells were analyzed. **D:** Scatter plot displays log_2_FC differential accessibility of on-target cell types compared to off target cells (x-axis) versus the on/off target expression ratio for CRE barcode expression (y-axis). On-target cell type definition and ratio are the same as C. Dashed line indicates the correlation between the on/ff target expression ratio and on-target accessibility of all 41 CREs (Spearman’s *r* = 0.26); solid line indicates the same correlation after removing CREs with viral barcodes detected only in off-target cell types (Spearman’s *r* = 0.47).

We next examined the extent to which subtype-specific chromatin accessibility predicts CRE-driven reporter expression across DRG neuronal subtypes. We calculated expected on/off target expression ratios for each CRE based on the pattern of differential chromatin accessibility at each CRE’s locus (see Methods; Table S15); CRE specificity ratios ranged from 0 to >100 for their respective on-target DRG neuron class. Overall, we observed a positive correlation (Spearman’s r = 0.26, p < 0.05) between the measured differential chromatin accessibility at each CRE locus and the CRE-driven reporter expression in each DRG cell type (Figure 4D), consistent with our attempt to prioritize functional CREs for *in vivo* testing. Interestingly, a subset of CREs in our multiplex screen exhibited high differential accessibility in their target cell types compared to their off target cell types, but their corresponding AAV barcodes were only detected in off target cell types (On/off target expression ratios = 0, > 6 cells). While our goal was to identify CREs that act as cell-type-specific enhancers, our findings suggest that some putative CREs might instead function as cell-type-specific silencers or require additional native chromatin context to enhance expression in their on-target cell types. Excluding these discordant CREs significantly improved the correlation between on target accessibility and on/off target AAV expression ratio from 0.26 to 0.47 (p < 0.05, Figure 4D).

Taken together, this multiplexed CRE scRNA-seq screen enabled the functional characterization of dozens of CREs in parallel as well as the discovery of several new enhancer AAVs capable of targeting distinct DRG neuronal subtypes.

### Modeling cis-regulatory logic of DRG neurons

The correlation of cell-type-specific chromatin accessibility with CRE activity in DRG neurons prompted us to ask if we could quantitatively model the *cis-*regulatory logic of DRG nociceptors. Previous studies have leveraged machine learning approaches such as convolutional neural networks (CNNs) to develop models trained on snATAC-seq data that can predict cell-type-specific chromatin accessibility based on sequence alone.^42–44^ We thus constructed a CNN, Predicting Accessibility In Nociceptors (PAIN-net), trained on DNA sequences from our mouse DRG snMultiome data, to predict both the neuronal subtype-specific chromatin accessibility signal intensity and the Log_2_FC of differential accessibility in each neuronal subtype compared to all other subtypes (see Methods). We tested model performance on randomly selected sequences comprising ∼15% of the mouse DRG snMultiome data that was excluded from training, and observed a median Pearson’s correlation of 0.72 across cell types between model-predicted and measured differential accessibility (Figure 5A; Figure S7A; Table S16). This model’s performance is comparable to chromatin accessibility predictive models generated in other tissues.^43^

**Figure 5:**
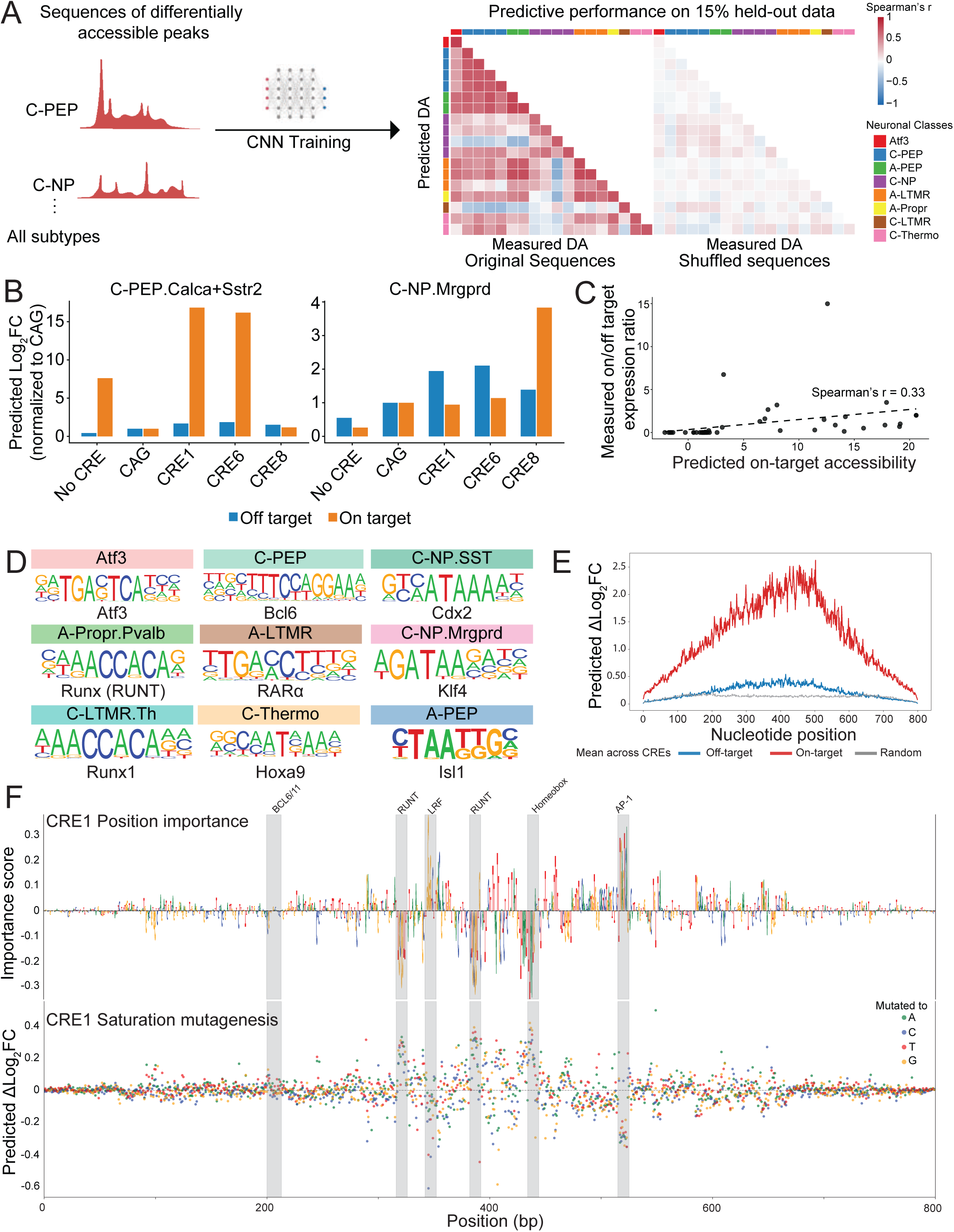
Machine learning prioritization of candidate CREs. **A:** (*Left*) Workflow schematic displays PAIN-net model training and validation. PAIN-net was trained on sequences of differentially accessible peaks (in each neuronal subtype compared to all others) to predict cell type-specific accessibility (log_2_FC in each neuronal subtype compared to all others) and peak signal. Training was performed on 85% of the data, while testing was performed on the held-out 15%. (*Right*) Heatmap displays the Pearson correlation between predicted and measured differential accessibility for each neuronal subtype (log_2_FC compared to all other neuronal subtypes). Results are shown for the held-out 15% data (median r = 0.72) and shuffled sequences (median r = -0.07). Color bars indicate broader neuronal classes and the order of subtypes within those classes is from the top to bottom: Red: Atf3, Blue: C-PEP (C-PEP.Calca.Adra2a, C-PEP.Calca.Dcn, C-PEP.Calca.Oprk1, C-PEP.Calca.Sstr2), Green: A-PEP (A-PEP.Calca.Smr2, A-PEP.Calca.Bmpr1b), Purple: C-NP (C-NP.Mrgpra3.Mrgprb4, C-NP.Mrgpra3.Trpv1, C-NP.Mrgprd, C-NP.Sst), Orange: A-LTMR (A-LTMR.Ntrk3^high^.Ntrk2, A-LTMR.Ntrk3^high^.S100a16, A-LTMR.Ntrk3^low^.Ntrk2), Yellow: A-Propr.Pvalb, Brown: C-LTMR.Th, Pink: C-Thermo (C-Thermo.Rxfp1, C-Thermo.Trpm8). **B:** PAIN-net recapitulates cell type specificity of CRE1, 6, and 8. Predicted log_2_FC was calculated for CRE1, CRE6, CRE8 and CAG control using PAIN-net for C-PEP.Calca+Sstr2 (*Left*) and C-NP.Mrgprd (*Right*) populations. All scores are normalized to the predicted log2FC of CAG. Predicted log2FC is defined as the PAIN-net predicted accessibility for a given neuronal subtype compared to all other neuronal subtypes. Orange bar (on-target): Predicted log2FC of C-PEP.Calca+Sstr2 (*left;* target subtype for CRE1 and CRE6) or C-NP.Mrgprd (*right;* target subtype for CRE8). Blue bar (off-target): Sum of predicted log2FC for all cell types aside from C-PEP.Calca+Sstr2 (*left*) or C-NP.Mrgprd (*right*). **C:** PAIN-net predictions correlate with CRE driven AAV barcode expression. Scatter plot shows PAIN-net predicted on-target differential accessibility versus the on/off target expression ratio of CRE barcode expression driven in the multiplexed AAV scRNA-seq screen (Spearman’s r = 0.33, p < 0.05). On-target cell types are defined as having accessible chromatin at the CRE locus (Table S15; see Methods). On-target differential accessibility is defined as the accessibility of on-target cell types compared to off target cell types. **D:** Sequence logos display motifs significantly enriched (q-value < 0.05) in the top PAIN-net predicted sequences for each neuronal subtype, relative to GC-matched backgrounds. Top sequences are defined as the 1,000 regions with the highest predicted differential accessibility (Log_2_FC) compared to all other cell types. Each motif is labeled with the best-matching transcription factor predicted to bind. Sequence logo size is based on nucleotide frequency. **E:** Plot displays the cumulative impact of *in silico* mutations across an 800bp sequence window on PAIN-net predicted accessibility, averaged over 43 CREs. Predicted ΔLog_2_FC (y-axis) is calculated as the sum of the absolute Δlog_2_FC values across on target cell types (red line) or off-target cell types (blue line). The gray line serves as a negative control derived from random DNA sequences. **F:** *In silico* saturation mutagenesis of CRE1 identifies key bases driving differential accessibility in C-PEP.Calca+Sstr2. Vertical grey bands mark putative transcription factor binding sites that coincide with high-impact regions. (*Top*) Attribution sequence logos illustrate the contribution of each nucleotide to C-PEP.*Calca*+*Sstr2* differential accessibility. Nucleotide height represents the predicted negative average impact of mutation on accessibility. (*Bottom*) Saturation mutagenesis scatter plot depicting the predicted impact (Δlog_2_FC compared to original sequence) of every possible single-nucleotide substitution. Points are colored by the substituted base.

A key motivation for developing PAIN-net was to prioritize CREs that drive cell-type-specific AAV expression in DRG. We thus asked whether PAIN-net-predicted CRE accessibility across DRG cell types is associated with each CRE’s *in vivo* specificity, as measured by their scRNA-seq on/off target expression ratio. PAIN-net correctly predicted the nociceptor subtypes in which CRE1, CRE6 and CRE8 were most active *in vivo*: C-PEP.Calca+Sstr2 (CRE1 and CRE6) and C-NP.Mrgprd (CRE8) (Figure 5B). We also observed good model performance across the *in vivo* multiplexed CRE screen; PAIN-net predictions were positively correlated with the measured on/off target expression ratios measured by scRNA-seq (Spearman’s r = 0.33; p-value <0.05; Figure 5C; correlation between predicted and measured accessibility in Figure S7B). These findings suggest that PAIN-net can be a valuable tool for nominating CREs likely to drive neuronal subtype-specific expression in the DRG.

We next asked which TF motifs are enriched in the sequences PAIN-net predicts to have the greatest differential accessibility in each DRG neuronal subtype compared to random sequences. PAIN-net identified TFs known to contribute to sensory neuron development (e.g. BCL6 in C-PEP nociceptors, RUNT family members in proprioceptors, RUNX1 in C-LTMR.Th neurons, and ATF3 in Atf3 neurons),^13,45,46^ as well as several new TFs that warrant further investigation (Figure 5D). These results are also in line with those observed in our SCENIC+ analysis of chromatin accessibility data (Figures 2F-G).

We next employed PAIN-net to assess the independent regulatory potential of key transcription factors by embedding their consensus binding motifs into random background sequences and predicting the resulting chromatin accessibility across neuronal subtypes. We found that *in silico* disruption of these embedded ATF3, RUNX1, and ISL1 consensus motifs significantly altered predicted accessibility (FDR < 0.01), whereas mutations in the flanking random regions yielded negligible effects (Figure S7C). Importantly, disruption of TF consensus motifs preferentially reduced chromatin accessibility in the specific neuronal subtypes in which the corresponding TF functions (e.g., ATF3 in the *Atf3* neurons) (Figure S7C). This indicates that PAIN-net has successfully captured cell-type-specific *cis-*regulatory dependencies of known motifs.

Having tested the model on known consensus sites, we next sought to map the precise nucleotides driving accessibility in the nociceptor CREs we screened *in vivo*. To do this, we performed *in silico* saturation mutagenesis using PAIN-net to systematically predict the impact of every possible single-nucleotide substitution on chromatin accessibility across each CRE. *In silico* mutagenesis revealed that the most impactful substitutions clustered at the center of the CRE sequences (Figure 5E), coinciding with the summits of differentially accessible peaks. Notably, the predicted magnitude of these substitutions was greater for a CRE’s on-target cell types than its off-target type (Figure 5E), suggesting that the model captures cell-type-specific regulatory logic. Applying this unbiased mutagenesis to CRE1 revealed that the specific nucleotides driving differential accessibility coincide with known transcription factor binding motifs, including those for BCL6/11, RUNT, Homeobox, and AP-1 (Figure 5F). Collectively, these findings indicate that PAIN-net captures the *cis*-regulatory logic used by DRG neuronal subtypes to establish unique gene expression patterns. This framework provides a powerful tool to both nominate native nociceptor CREs for testing *in vivo* and to design synthetic elements with the potential to achieve greater cell-type-specificity than is observed with natural CREs.^31,35,36^

### Functional and translational applications of nociceptor-specific AAVs

Given the long-term goal of developing cell-type-specific AAVs to inhibit nociceptor activity, we next generated AAVs in which either the nociceptor-specific CRE1 or the pan-expressing CAG promoter drives expression of the inward-rectifying potassium channel K_ir_2.1 and mCherry (Figure 6A). We also constructed a control AAV using the CAG promoter to drive expression of mCherry alone. AAVs were packaged in the AAV-PHP.S capsid^37^ and delivered *in vitro* to dissociated DRG cells cultured for whole cell patch clamp electrophysiology. CAG-K_ir_2.1- and CAG-mCherry-transduced neurons were identified by bright mCherry fluorescence and CAG-K_ir_2.1 neurons exhibited significantly reduced excitability compared to mCherry-transduced neurons as measured by an increase in the amount of injected current required to stimulate action-potential firing (rheobase) (Figures 6B-C). Expression of mCherry in CRE1-K_ir_2.1-transduced neurons was too dim to reliably distinguish all cells from autofluorescence, but we observed a population of small-to-medium diameter neurons^47,48^ with rheobase currents higher than any measured in mCherry control cultures (p < 0.0001, One-way ANOVA; Figures 6B-C; Figure S8A; see Methods). Both CAG-K_ir_2.1 and high rheobase CRE1-K_ir_2.1 neurons demonstrated hyperpolarized membrane potentials (p < 0.0001, One-way ANOVA; Figure 6D), and larger inward-rectifying currents at hyperpolarized potentials (Figure 6E). The increased potassium conductance at sub-threshold voltages likely underlies this observed hyperpolarization in the resting membrane potential and is consistent with the known hyperpolarizing effects of K_ir_2.1 overexpression.^49^ Collectively, these data demonstrate that CRE1-driven K_ir_2.1 expression reduces DRG neuronal excitability.

**Figure 6:**
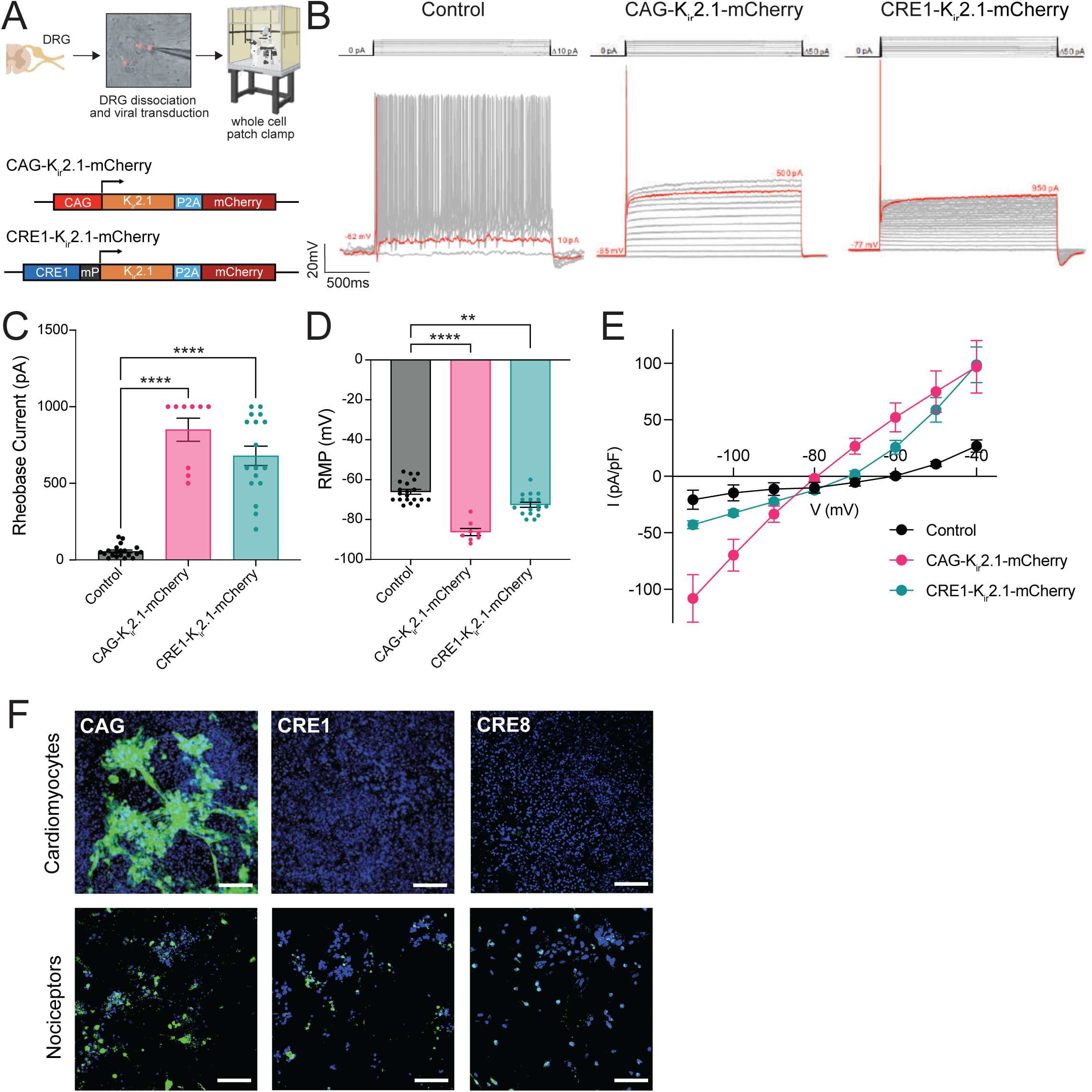
Functional and translational applications of nociceptor-specific AAVs. **A:** (*Top*) Workflow for DRG dissociation, viral transduction, and whole-cell patch-clamp recording. (*Top, Middle*) Representative image of CAG infected cultured DRG neurons (red). (*Bottom*) Construct maps for the experimental vectors expressing K_ir_2.1 under the ubiquitous CAG promoter or CRE1 and mP (minimal promoter). **B:** Representative voltage responses to current injection steps (step protocol shown above) from Control (*Left*), CAG-K_ir_2.1-mCherry (*Middle*), and CRE1-K_ir_2.1-mCherry (*Right*) cells. Control neurons exhibit repetitive firing. In contrast, CAG and CRE1 neurons exhibit a failure to fire at low current steps, with red traces indicating the increased rheobase. **C:** Summary data showing the minimum current required to elicit an action potential (rheobase). Neurons expressing K_ir_2.1 via CAG and CRE1 showed significantly higher rheobase compared to controls (Control, n = 20; CAG, n = 9; CRE1, n = 17). Data are presented as mean ± SEM. One-way ANOVA with multiple comparisons (* p < 0.05, ** p < 0.01, **** p < 0.0001) (F(2, 43) = 76.72, p < 0.0001). Post hoc analysis using Dunnett’s multiple comparisons test revealed that rheobase was significantly increased in both the CAG group (p < 0.0001) and the CRE1 group (p < 0.0001) compared to control. Sample sizes (n) represent individual neurons recorded across at least 2 independent preparations. **D:** Quantification of resting membrane potential (RMP) showing a significant hyperpolarization of neurons expressing K_ir_2.1 both via CAG and CRE1 compared to control cells (Control, n = 20; CAG, n = 8; CRE1, n = 17). Data are presented as mean ± SEM. One-way ANOVA with multiple comparisons (* p < 0.05, ** p < 0.01, **** p < 0.0001) (F(2, 42) = 40.08, p < 0.0001). Post hoc analysis using Dunnett’s multiple comparisons test showed that RMP was significantly hyperpolarized in the CAG group compared to control (p < 0.0001), as well as in the CRE1 group (p = 0.0014). Sample sizes (n) represent individual neurons recorded across at least 2 independent preparations. **E:** Current-voltage (I-V) relationships for cells in Control, CAG-K_ir_2.1-mCherry, and CRE1-K_ir_2.1-mCherry groups. Steady-state current density (pA/pF) plotted against membrane potential (mV). Neurons expressing K_ir_2.1 via CAG and CRE1 exhibit characteristic inward-rectifying currents at hyperpolarized potentials compared to Control cells, consistent with expression of K_ir_2.1 (Control, n = 6; CAG, n = 6; CRE1, n = 11). **F:** Representative immunofluorescence images of iPSC-derived cardiomyocytes and nociceptors. Cells were transduced with AAV vectors expressing EGFP driven by the CAG promoter, CRE1, or CRE8. Images are representative of n = 3-5 independent experiments. Blue = DAPI; Green = EGFP. Scale bars = 100 µm.

To assess whether our nociceptor CREs are likely to drive AAV gene expression preferentially in human nociceptors, we cultured human iPSCs (LiPSC-GR1.1) and differentiated them using protocols that produce either nociceptor-like sensory neurons or cardiomyocytes.^50,51^ We transduced these cells with AAV-PHP.S vectors expressing an eGFP reporter under the control of CRE1, CRE8, or the ubiquitous CAG promoter. After 3-4 weeks in culture for functional maturation, we assessed reporter expression by immunofluorescence microscopy. We observed robust reporter expression driven by the CAG promoter in both iPS nociceptors and cardiomyocytes, but CRE1 and CRE8 only drive reporter expression in nociceptors (Figure 6F). These findings support the possibility that nociceptor CREs identified and screened in mice can function similarly in nociceptors across species.

## Discussion

Peripheral sensory neurons comprise diverse subtypes that collectively encode touch, temperature, itch, and pain. Chronic pain is linked to nociceptor hyperexcitability, yet translating mechanistic insight into precise therapeutics has been constrained by the lack of subtype-selective access across species.^52,53^ Here we built matched single nucleus multi-omic atlases of mouse and human DRG, defined conserved cell-type-specific regulatory programs, and identified CREs that drive gene expression preferentially in nociceptor subtypes *in vivo* and in human iPSC-derived nociceptor-like neurons. We also developed a sequence-based model (PAIN-net) that captures key components of nociceptor *cis*-regulatory logic, enabling both prioritization of CREs likely to function as enhancers *in vivo* and design of synthetic elements with tunable properties. Together, these resources (available at www.painseq.com) support a generalizable strategy for cell-type-specific neuromodulation in the peripheral nervous system.

Enhancer AAV tool development relies on cell-type–resolved epigenomic maps; in the CNS, such maps have enabled nomination of CREs that function *in vivo* when placed upstream of minimal promoters in AAV cassettes.^14–17^ By generating matched mouse and human DRG snMultiome atlases, we extend this capability to the peripheral nervous system, where neuronal subtype diversity is well described transcriptionally but has remained difficult to access genetically without transgenic mouse lines. The overlap we observe between mouse and human DRG regulatory features provides a mechanistic rationale for using mouse *in vivo* screens as a discovery engine for elements likely to function in human nociceptors, while also establishing a reference for interpreting species-specific divergence that may influence translation.

Our multi-omic analyses extend prior transcriptomic descriptions of DRG^5,6,30–32^ by defining subtype-resolved *cis*-regulatory landscapes and TF-centered regulatory networks. Gene regulatory network analyses highlight candidate regulators across sensory lineages (e.g. enrichment of RUNX-family activity in proprioceptors) and link these programs to specific putative enhancers. This framework supports CRE nomination using convergent evidence, including subtype-enriched accessibility, cross-species sequence conservation, and membership in TF-centered regulatory modules. As chromatin accessibility alone does not guarantee functional enhancer activity *in vivo,* this multi-criterion prioritization framework helps address an important bottleneck in enhancer engineering.^56,57^ That said, CRE prioritization remains imperfect because selectivity and expression level can change substantially when CREs are placed in an AAV cassette context. Accordingly, functional AAV screens of prioritized CREs using quantitative, cell-type–resolved readouts remain essential.

Our functional AAV screen in mouse DRG identified a subset of nominated CREs that drive reporter expression preferentially in nociceptor classes, including enhancers biased toward peptidergic (CRE1, CRE6) and non-peptidergic populations (CRE8). These tools were successfully validated using both FISH to assay co-localization with canonical markers and scRNA-seq to quantify full subtype-resolved specificity. While three individually tested CREs showed specificity for their intended nociceptor classes, our multiplexed barcoded CRE screen enabled us to functionally rank dozens of elements within the same AAV library. This approach identified high-performing nociceptor CREs (e.g., CRE114) as well as elements that preferentially drive expression in non-nociceptive DRG populations (e.g., thermosensory, proprioceptive, or mechanosensory lineages).

The multiplexed screen also provided mechanistic insight into how endogenous chromatin accessibility relates to cell-type expression patterns driven by an AAV. The modest but significant correlation between subtype-enriched chromatin accessibility and *in vivo* barcode expression supports accessibility as a useful guide for enhancer discovery while underscoring the contribution of additional regulatory layers. For example, some CREs showed strong on-target enrichment, whereas other elements with similar accessibility profiles produced off-target expression. We also observed a subset of highly accessible CREs that drove expression only in off-target cells, suggesting that these sequences could act as context-dependent silencers. Together, these observations motivate continued empirical screening and the development of predictive models that can both prioritize candidate CREs and guide the design of synthetic elements that improve specificity, strength, and/or cross-species portability.

To this end, we developed PAIN-net, a sequence-based model that predicts neuronal subtype–specific chromatin accessibility directly from DNA sequence and provides a quantitative framework for interpreting enhancer “grammar.” PAIN-net generalizes to held-out loci and tracks measured CRE-driven expression patterns, supporting the view that a substantial component of nociceptor *cis*-regulatory logic is learnable from sequence features. Motif enrichment and *in silico* perturbation analyses further suggest that PAIN-net captures biologically meaningful dependencies on transcription factor binding sites and their arrangement. More broadly, PAIN-net brings model-guided CRE design into the peripheral sensory system and establishes a foundation for optimizing CRE specificity and strength or for combinatorial targeting strategies. In practice, such design capabilities could enable viral tools tailored to distinct experimental or therapeutic goals, including broader or more selective targeting of nociceptor subtypes, or exclusion of specific proprioceptive and mechanosensory neurons to minimize sensory side effects.

Consistent with the potential of nociceptor-biased AAVs as a future pain therapeutic, CRE1 drove sufficient expression of K_ir_2.1 to reduce DRG neuron excitability *in vitro*, with electrophysiological signatures consistent with increased inward-rectifying potassium conductance and hyperpolarization. This result demonstrates that even comparatively modest enhancer-driven expression can yield meaningful functional effects when paired with a potent effector. It also highlights a design principle relevant to therapeutic translation: specificity and expression strength can often trade off. For certain effectors, sub-maximal expression may still provide robust modulation while reducing over-expression toxicity and provide a wider window to fine-tune expression.^58^ The robust and specific expression of CRE1 in human iPSC-derived nociceptors over cardiomyocytes supports the translational potential that conserved *cis*-regulatory logic can be leveraged to target human nociceptors.

Several limitations define priorities for future work. First, human neuronal chromatin accessibility data were comparatively sparse, necessitating broader grouping of neuronal subtypes than in mice for portions of the epigenomic analysis. Expanding high-quality human multi-omic sampling across anatomical levels, ages, injury states, and diverse donors will sharpen enhancer nomination and help identify regulatory programs that are human-specific or state-dependent. Second, the diversity and scarcity of sensory neurons in peripheral ganglia fundamentally limits CRE screening throughput. Development of nociceptor iPSC models that better recapitulate *in vivo* nociceptor subtypes could enable ultra-high throughput synthetic enhancer screens that are currently being performed in cell lines in other contexts.^42^ Third, our *in vivo* AAV screening used neonatal ICV delivery and a viral capsid optimized for broad DRG transduction in mice. Translation will require validating enhancer performance across multiple delivery routes, capsids, and evaluation of off-target expression across multiple tissues.

Despite these limitations, the cross-species DRG multi-omic atlases, predictive models, and nociceptor-selective AAVs presented here provide important first steps toward a modular toolkit for peripheral neuromodulation across species. This framework should extend to other peripheral ganglia and autonomic circuits where genetic access remains limited.

## Methods

### Animals

All mouse experiments in this study were approved by the National Institutes of Health and the Harvard Medical School IACUC. Experiments followed the ethical guidelines outlined in the NIH Guide for the Care and Use of Laboratory Animals. Mice were housed under standard conditions, with no more than five animals per cage, maintained on a 12 h day/night cycle, with food and water. Both male and female mice were used for experiments, aged between 6-24 weeks, and the minimum number of animals was used for each experimental group. Animals were maintained on a C57BL/6J genetic background.

### Mouse nuclear isolation

Dorsal root ganglia from all spinal levels were dissected from 50 mice (males and females) aged 6 to 24 weeks. All DRGs were directly frozen on dry ice and stored at -80C until processing for single-nucleus RNA sequencing (snRNA-seq), using a detergent-based nuclei isolation protocol optimized for neuronal tissue. All reagents were prepared on the same day and kept ice-cold unless otherwise specified. RNase-free technique was used throughout.

On the day of preparation, a swinging-bucket centrifuge was pre-cooled to 4°C. Nuclei lysis buffer (NST), resuspension buffer (ST), and 1× nuclei buffer (10x Genomics) were prepared fresh, supplemented with RNase inhibitor (RNasin Plus, Promega N2611), and maintained on ice. Dounce homogenizers and Tissue Tearor probes were thoroughly cleaned between samples using nuclease-free water and 70% ethanol to prevent cross-contamination. NST (lysis) buffer consisted of 1×stock buffer (20 mM NaCl, 20 mM Tris-HCl pH 7.5, 1 mM CaCl₂, 6 mM MgCl₂), 0.2% IGEPAL CA-630 (Sigma I8896-50ML), and 0.84% BSA in nuclease-free water. ST buffer consisted of 1×stock buffer supplemented with 0.9% BSA. All buffers were vacuum-filtered, stored at 4°C, and supplemented with RNase Inhibitor immediately prior to use.

Pooled frozen DRGs from 5 mice were processed each time. Tissue was transferred from dry ice and briefly warmed at the tube edges to facilitate homogenization. Samples were lysed in NST buffer containing IGEPAL CA-630 and BSA, supplemented with RNase inhibitor. Tissue was mechanically disrupted using a Tissue Tearor on a low-speed setting (2–3) for 5 seconds, ensuring complete coverage of the tissue. Homogenates were kept on ice immediately following disruption. The homogenate was transferred to a glass Dounce homogenizer and gently washed to recover residual tissue. Samples were further homogenized using a tight pestle with 15 slow strokes, avoiding bubble formation. The resulting homogenate was passed through a 40 µm cell strainer into a pre-chilled 15 mL conical tube, followed by an additional wash of the Dounce homogenizer with ST buffer to maximize nuclei recovery. Filtered homogenates were centrifuged at 500×g for 10 minutes at 4°C. Supernatant was carefully removed, leaving approximately 0.5 mL above the pellet. The nuclei pellet was gently resuspended in 1× nuclei buffer (10x Genomics, PN-2000207) as needed.

Nuclei suspensions were transferred for fluorescence-activated nuclei sorting (FANS) performed by the BWH Center for Cellular Profiling. Collection tubes were pre-coated with 1× nuclei buffer to reduce nuclei loss. Sorting was performed using a 70 µm nozzle. Nuclei were gated based on GFP signal (neuronal nuclei are GFP+) at forward and side scatter, exclusion of debris, and DNA staining using 7-aminoactinomycin D (7-AAD; final concentration 10 ng/µL). Negative controls were used to establish gating parameters.

Sorted nuclei were collected directly into tubes containing 100ul of nuclei buffer. To concentrate nuclei in 5ul 1× nuclei buffer for downstream 10x Genomics multiome application, concanavalin A-conjugated paramagnetic beads (CUTANA) were used as previously described.^59^ For snRNA-seq experiments, nuclei were taken directly into library preparation without additional purification or centrifugation to minimize loss.

Nuclei concentration was determined using a hemocytometer following dilution with Trypan blue or DAPI staining. Counts were averaged across four quadrants and used to calculate nuclei concentration (nuclei/µL). An aliquot of nuclei was optionally inspected by fluorescence microscopy to confirm nuclear integrity and morphology.

### Mouse snMultiome-seq library preparation and sequencing

Nuclei suspensions were concentrated and approximately 10,000 nuclei per library were loaded on the 10x Genomics Chromium Controller, following the manufacturer’s guidelines for the Chromium Next GEM Single Cell Multiome ATAC + Gene Expression kit. Library preparation, GEM generation, and barcoding were performed according to the manufacturer’s protocol.

### Mouse multi-ome data processing

#### Read alignment and ambient RNA correction

Raw reads collected from mice DRG samples were aligned to the Mouse mm10 (GENCODE vM23/ensembl98) mouse reference genome using the Cell Ranger-ARC pipeline (v2.0.0; default settings). The reference genome was downloaded from the 10x Genomics website. To ensure high-quality data integration, RNA counts were filtered for ambient noise using CellBender^60^ (default settings).

#### RNA modality

Seurat v5 was used to process the RNA modality by combining all preprocessed in-house and published ambient RNA corrected count matrices (18 libraries total). Nuclei with more than 1,000 detected genes and less than 25% mitochondrial counts were retained. Filtered data were normalized using log-normalization, and the top 2,000 variable genes were identified, excluding mitochondrial genes. Data were then scaled, and principal component analysis (PCA) was performed using the selected variable genes. Batch correction and integration across libraries were performed using the Seurat CCA method. The number of principal components (PCs) was determined as the point where selected PCs cumulatively explained at least 90% of the variance and additional PCs explained less than 5% of the standard deviation of highly variable gene expression. We then clustered the data at a resolution of 1 and clusters were embedded in UMAP space. Clusters were then annotated based on the enrichment (Log_2_FC > 0.5, adjusted p-value < 0.05 relative to all other clusters) of the following marker genes: *Snap25*, *Rbfox3*, and *Pvalb,* as neurons; *Fabp7* and *Ednrb* as for SGCs; *Mpz* and *Mbp* for mySCs; *Scn7a* for nmSC; *Ptprc and Ccr2*for Immune cells; *Pecam1* and *Flt1* for Endothelial cells; *Notch3* and *Kcnj8* for Pericytes; *Pdgfra* and *Tbx18* for Fibroblasts. Clusters showing enrichment of both neuronal and non-neuronal markers were annotated as doublets and removed.

Neurons were then subsetted and re-clustered. For this step, the QC threshold was increased to retain nuclei with more than 1,500 detected genes and less than 15% mitochondrial gene counts. We then predicted neuronal subtype identity from the mouse harmonized DRG atlas^6^ using Seurat’s label transfer method, and excluded nuclei with a prediction score less than 0.5.

With the exception of three mouse libraries, all libraries had a mixture of male and female mice and thus we performed an additional computational step in assigning sex labels to each nucleus. Sex labels were assigned to individual nuclei based on expression of the Y-chromosome gene *Uty* and the X-inactivation transcript *Xist*. Cells were classified as “Male” if *Uty* expression exceeded the 75th percentile while *Xist* remained below the 75th percentile, and as “Female” if *Xist* exceeded the 75th percentile while *Uty* remained below the 75th percentile; cells not meeting these criteria were initially labeled as “Ambiguous”. For the three libraries where only one sex was sequenced, sex labels were manually assigned (mDRG_rest_day3, Techameena_ctrl_F, and Techameena_ctrl_M).

### ATAC modality

Signac (v1.16.0) was used to process the ATAC modality. Peak files (.bed) and fragment files generated by Cell Ranger ARC were used to construct peak-by-cell count matrices for each of the 18 libraries, which were then merged into a single object. ATAC data were subsetted to retain only RNA-matched cells, and RNA-derived cell identities were transferred to the ATAC modality based on shared barcodes. Nuclei with fewer than 1,000 total fragments in peaks or TSS enrichment scores below 3 were excluded. Cell-type specific peaks were called using MACS2 in BEDPE mode, restricted to standard chromosomes and excluding blacklist regions. The resulting peak set was used for all downstream analyses. This workflow was applied both to the full set of annotated cell types (neuronal and non-neuronal populations) and to a neuron-focused analysis in which neuronal subtypes were considered separately.

### Pseudobulk-based differential accessibility of mouse DRG cell types for CRE prioritization

To identify differentially accessible (DA) regions in mouse DRG cell types, we used a pseudobulk strategy implemented in Seurat/Signac and DESeq2. Briefly, cells were grouped by annotated cell type and randomly assigned to one of *n* = 10 synthetic pseudoreplicates per cell type (set.seed = 42 for reproducibility). For each cell type-pseudoreplicate combination, raw accessibility counts from the MACS2-called peak matrix were aggregated using AggregateExpression to generate a pseudobulk count matrix with one column per pseudoreplicate.

For DA testing, we performed repeated one-vs-rest comparisons. A model was fit with design ∼ celltype_replicateID. Peaks with low overall signal were removed prior to modeling (retaining features with total counts ≥ 10 across all pseudobulk samples). Size factors were estimated using DESeq2’s poscounts method to accommodate sparsity typical of accessibility data, and the negative binomial model was fitted using fitType = “local.” Peaks were considered differentially accessible based on adjusted p-value thresholds (FDR < 0.05 for all analyses) and effect size cutoffs (log_2_FC > 0.5), as reported.

#### Gene regulatory network analysis

To infer cell type-specific gene regulatory networks for the mouse DRG, we used SCENIC+ which required two inputs: (1) the pseudobulk-based DA regions in mouse DRG cell types, (2) a raw gene expression matrix of annotated mouse nuclei and (3) the binarized topic-region associations outputted from pycisTopic (method details below).

PycisTopic^24^ was used to group peaks into groups of regions with a common accessibility profile across cells (topics). Briefly, peak accessibility profiles were exported from a Signac object as a sparse peak-by-cell count matrix (MatrixMarket format) together with matching cell barcodes, peak coordinates, and per-cell metadata. These inputs were used to construct a cisTopic object, while filtering regions overlapping the ENCODE blacklist. Cell-level annotations were then added to the cisTopic object to enable downstream visualization and interpretation.

Topic modelling was performed using latent Dirichlet allocation (LDA) implemented in pycisTopic with MALLET. Multiple models were trained across a range of topic numbers (10–200 topics), using default parameters (seed=555) and multi-core parallelization (60 CPUs). Trained models were subsequently evaluated using standard topic model selection metrics (Arun_2010, Cao_Juan_2009, Mimno_2011, and model log-likelihood). Based on these diagnostics, a model with 48 topics was selected for major DRG cell classes, and for DRG neuronal subtypes 120 topics were selected. Topics were binarized to derive discrete topic-associated regulatory region sets (“candidate enhancers”), using two complementary approaches: Otsu thresholding and a rank-based strategy selecting the top 3,000 regions per topic. Binarized topic-region associations were exported as sorted BED files as the input for SCENIC+.

For the pycisTopic and SCENIC+ analysis of the mouse multi-ome data across major cell classes (neuronal and non-neuronal populations), the dataset was first randomly downsampled to a maximum of 1,000 nuclei per cell type. For the pycisTopic and SCENIC+ analysis of neuronal DRG subtypes, the full neuronal dataset was used.

### Human DRG acquisition and dissection

All procedures for human tissue procurement and ethical oversight were reviewed and approved by the Institutional Review Board at Brigham and Women’s Hospital [2024P002201/MGB1924]. Protocol IRB#2017P000757 was reviewed by Mass General Brigham IRB and determined to be not human research. DRGs were collected from consented donors using a rapid autopsy protocol at Mass General Brigham as described in Bhuiyan et al., 2024.^6^ All donors were deidentified. Age and sex of donors are reported in Table S1.

### Human nuclear isolation

Single nuclei of human DRGs were collected using a previously described gradient protocol at Harvard Medical School.^61^ Briefly, human DRGs were initially pulverized on dry ice and stored at -80C until processing. Approximately pulverized tissue from 4 donors was used per experiment. and ∼0.5 to 1 cm^3^ of powder was placed into homogenization buffer [0.25 M sucrose, 25 mM KCl, 5 ml of MgCl_2_, 20 mM tricine-potassium hydroxide (KOH; pH 7.8), 1 mM dithiothreitol (DTT), actinomycin (5 μg/ml), 0.04% BSA, and ribonuclease (Rnase) inhibitor (0.1 U/μl)] for ∼15 seconds on ice. After the brief incubation for 15 seconds on ice, samples were transferred to a Dounce homogenizer for an additional 10 strokes with a tight pestle in a total volume of 5 mL of homogenization buffer. IGEPAL was added to a final concentration of 0.32%, and five additional strokes were performed with the tight pestle. The tissue homogenate was then passed through a 40-μm filter and diluted 1:1 with a working solution. Neuronal nuclei were enriched as previously described^20^ and concentrated using concanavalin A-conjugated paramagnetic beads (CUTANA).^59^

### Human snMultiome-seq library preparation and sequencing

Nuclei were labeled with DAPI to inspect their integrity under a fluorescent microscope and determine their quantity manually using a hemocytometer. Approximately, 10,000 nuclei were used for each ATAC and gene expression library. Libraries were generated according to the Chromium Next GEM Single-cell Multiome ATAC + Gene Expression guidelines (10x Genomics). Both ATAC and RNA libraries were shipped to GENEWIZ (Azenta Life Sciences) and were sequenced based on the sequencing parameters suggested by 10x Genomics on the Illumina NovaSeq X Plus platform to an approximate read depth of ∼25,000 reads per cell (multiome) and ∼20,000 reads per cell (RNA).

### Human multi-ome data processing

#### Read alignment and ambient RNA correction

Raw reads from the human samples were aligned to the Human GRCh38 (GENCODE v32/Ensembl98) using the Cell Ranger-ARC pipeline (v2.0.0; default settings). The reference genome was downloaded from the 10x Genomics website. To ensure high-quality data integration, RNA counts were filtered for ambient noise using CellBender^60^ (default settings).

#### Genotyping and demultiplexing

To maximize cost and efficiency, snRNA-seq was performed on batches of 1-13 donors. We used Souporcell to demultiplex sequencing reads to the level of individual donors. Briefly, Souporcell detects genetic variants within sequenced RNA, generates a cell by variant genotype matrix, and clusters cells based on their allelic assignments.^62^ Souporcell version 2.5 was run using default settings. Clusters were initialized with known genotypes (imputed from DNA sequencing) provided by the -known_genotypes option. As input, we combined single cell RNA runs containing the same donors. We verified that samples contained the expected donors by cross-correlating cluster genotypes for each sample against imputed genotypes for all donors. We restricted analysis to Souporcell clusters whose genotypes were correlated with imputed genotypes with a Pearson correlation coefficient of greater than 0.7.

For genotyping, DNA purification was performed with up to 25mg of tissue using the QIAamp Fast DNA TissueKit and sent to Azenta Life Sciences for 4X WGS. Imputation was performed using GLIMPSE2 and the HGDP-1kGP reference panel. Only common variants (minor allele frequency > 5%) were used for Souporcell cluster initialization.

#### RNA modality

Seurat v5 was used to process the RNA modality by combining all ambient RNA corrected count matrices (33 libraries total). Nuclei with more than 500 detected genes and less than 20% mitochondrial counts were retained. Filtered data were normalized using log-normalization, and the top 2,000 variable genes were identified. Data were then scaled, and PCA was performed using the selected variable genes. Batch correction and integration across libraries were performed using the Seurat CCA method. The number of PCs was determined as the point where selected PCs cumulatively explained at least 90% of the variance and additional PCs explained less than 5% of the standard deviation of highly variable gene expression. We then clustered the data at a resolution of 0.7 and clusters were embedded in UMAP space.

Clusters were then annotated based on the enrichment (Log_2_FC > 0.5, adjusted p-value < 0.05 relative to all other clusters) of the following marker genes: *SNAP25*, *PRPH*, and *NEFH*, as neurons; and *SPARC*, *MPZ*, and *PTPRC*, as non-neuronal populations. Clusters showing enrichment of both neuronal and non-neuronal markers were annotated as doublets and removed. Non-neuronal clusters were then further annotated using *FABP7* and *EDNRB* as markers for SGCs; *MPZ* and *MBP* for mySCs; *SCN7A* for nmSC; *PTPRC*, *CCR2* and *CD3E* for Immune cells; *PECAM1* and *FLT1* for Endothelial cells; *NOTCH3* and *KCNJ8* for Pericytes; and *PDGFRA* and *TBX18* for Fibroblasts.

Neurons were then subsetted for focused subtype analysis. We then predicted neuronal subtype identity using Seurat’s label transfer method by anchoring the neuronal subset to a human DRG atlas.^5^ For this step, anchor features were defined as genes shared between the dataset and the atlas, after removing non-coding RNAs and unknown genes. Nuclei with prediction scores greater than 0.5 were retained for downstream analysis.

To assign nuclei to respective human donors, Souporcell^62^ was used in genotype aware mode for all samples, except H32_NeuN for which genotype sequencing data was not available.

#### ATAC modality

For quality control, data were filtered using ArchR^63^ (v1.0.3) (number of fragments per cell > 1000, TSS Enrichment > 3). Signac (v1.16.0) was used to process the ATAC modality. Peak files (.bed) and fragment files generated by Cell Ranger ARC were used to construct peak-by-cell count matrices for each of the 18 libraries, which were then merged into a single object. ATAC data were subsetted to retain only RNA-matched cells, and RNA-derived cell identities were transferred to the ATAC modality based on shared barcodes. Cell-type-specific peaks were called using MACS2 in BEDPE mode, restricted to standard chromosomes and excluding blacklist regions. For neurons, the MACS2 peak calling was done with the “broad” parameter set to TRUE due to higher sparsity of the data. The resulting peak set was used for all downstream analyses. This workflow was applied both to the full set of annotated cell types (neuronal and non-neuronal populations) and to a neuron-focused analysis in which neuronal subtypes were considered separately.

#### Pseudobulk-based differential accessibility of human DRG cell types for CRE prioritization

Differential chromatin accessibility was assessed on a per–cell type basis using a pseudobulk strategy implemented in Seurat/Signac and DESeq2. Briefly, cells were grouped by annotated cell type and randomly assigned to one of *n* = 10 synthetic pseudoreplicates per cell type (set.seed = 42 for reproducibility). For each cell type–pseudoreplicate combination, raw accessibility counts from the MACS2-called peak matrix were aggregated using AggregateExpression to generate a pseudobulk count matrix with one column per pseudoreplicate.

For DA testing, we performed repeated one-vs-rest comparisons. A DESeq2 model was fit with design ∼ celltype_replicateID. Peaks with low overall signal were removed prior to modeling (retaining features with total counts ≥ 10 across all pseudobulk samples). Size factors were estimated using DESeq2’s poscounts method to accommodate sparsity typical of accessibility data, and the negative binomial model was fitted using fitType = “local”. Peaks were considered differentially accessible based on adjusted p-value thresholds (FDR < 0.05 for analyses for DRG cell types; FDR < 0.1 for DRG neuronal subtypes) and effect size cutoffs (log2FC > 0.5), as reported.

#### Gene regulatory network analysis

To infer cell type-specific gene regulatory network for the human DRG, we used SCENIC+, which required three inputs: (1) the pseudobulk-based DA regions in human DRG cell types, (2) the raw gene expression matrix of annotated human nuclei, and (3) the binarized topic-region associations outputted from pycisTopic. For pycisTopic, we used the same protocol as we did with mice, with the exception of using the ENCODE black list for humans and no downsampling was performed for any analyses.

### Differential gene expression across sexes

To identify sex-specific transcriptomic differences, we performed pseudobulk differential expression analysis using the Seurat and DESeq2 packages in R. We aggregated single-cell RNA counts by sample, sex, and cell type using AggregateExpression to create pseudobulk profiles, subsequently extracting metadata to reconstruct the experimental design. For each cell type with sufficient replication (minimum 2 replicates per sex), we performed differential expression analysis between male and female samples using FindMarkers with the DESeq2 test (test.use = “DESeq2”). To visualize the results, we generated volcano plots using ggplot2 and ggrepel, identifying significant genes (adjusted p-value < 0.05 and |log_2_FC| > 0.5).

### CRE prioritization

To prioritize candidate CREs for functional screening, we first identified genomic regions exhibiting significant differential chromatin accessibility in target neuronal subtypes (e.g., peptidergic or non-peptidergic nociceptors) compared to all other DRG cell types (Log_2_FC > 0.5, adj. p-value < 0.05). These candidate peaks were further filtered for evolutionary conservation by mapping mouse sequences to the human genome using the liftOver tool, ensuring the selection of elements with conserved regulatory potential. We additionally prioritized CREs predicted to function as active enhancers by SCENIC+, specifically those linked to cell-type-specific eRegulons or containing binding motifs for key lineage-defining transcription factors (e.g., RUNX1, BCL11A). Final candidate enhancer sequences were confirmed using custom tracks visualized via the UCSC Genome Browser. Genomic coordinates for peaks of interest (e.g., mm10) were located, and DNA sequences were extracted directly from the browser. For optimal cloning efficiency, enhancers were selected based on a size constraint of less than 1 kb in length.

### Construction of enhancer-driven AAV expression vectors

Enhancer sequences (see Table S17) were identified and analyzed for unique restriction sites to facilitate directional cloning. Enhancer templates were generated via genomic DNA extraction (QIAGEN 69504) or as synthetic gBlocks (IDT). Templates were amplified using Q5 High-Fidelity DNA Polymerase (NEB M0492S) under standard cycling conditions, verified by agarose gel electrophoresis, and purified (QIAquick, QIAGEN 28104).

Both the AAV backbone and enhancer inserts were digested with appropriate restriction enzymes (e.g., *MluI*, *SalI-HF*, or *XhoI*). To minimize self-ligation, vectors were dephosphorylated with Antarctic Phosphatase (NEB M0289S). Digested products were gel-purified and ligated using T4 DNA Ligase (NEB M0202S) at a 1:5 vector-to-insert molar ratio. Following transformation into *E. coli* (DH5⍺) (NEB C2987H), successful clones were screened via colony PCR or restriction analysis and validated by Sanger or whole-plasmid sequencing (Azenta Life Sciences). Validated constructs were produced using endotoxin-free Maxiprep (Thermo Fisher A31231) and submitted for AAV packaging.

### Multiplexed CRE library design

A multiplexed library was constructed containing 106 prioritized CREs, a mini CMV enhancer control, and a No CRE control (see Table S17). For each element, a 275bp sequence flanking the CRE summit was synthesized as an oPool (IDT) with the following architecture: 5’ InFusion overhang - CRE - restriction sites (SpeI, FseI) - barcode - 3’ InFusion overhang Unique barcodes were assigned using the R package DNAbarcodes, following the logic described in Hrvatin et al. 2019.^41^ The oPool was PCR-amplified and cloned into the AAV backbone as previously described.^64^ A functional reporter cassette (containing a minimal CMV promoter, mGreenLantern-KASH, and WPRE) was subsequently inserted into the library restriction sites (*SpeI/FseI*) as described^41^ with the following architecture:

SpeI–minimal CMV promoter--mGreenLantern-KASH--WPRE–FseI

The structure of the final AAV library is:

AAV ITR-CRE–minimal promoter–mGreenLantern-KASH–WPRE-barcode-polyA-AAV ITR

### Amplification of CRE library and barcoding

The CRE library was generated with an approach similar to that described in Hrvatin et al. 2019.^41^ In brief, the CRE library was generated by PCR amplification of a synthetic oPool (IDT) using Q5 High-Fidelity Master Mix (NEB). Amplicons (approx. 350 bp) were purified and verified via agarose gel electrophoresis. The AAV backbone was linearized through double restriction digestion and gel-purified to remove template DNA.

The purified oPool inserts were cloned into the linearized vector using In-Fusion Snap Assembly (Takara Bio 638947). To ensure high transformation efficiency, the assembly products were concentrated and desalted prior to electroporation into MegaX DH10B T1R cells (Invitrogen C6400-03). To maintain library representation, we ensured >100X coverage by plating the library on 245-mm BioAssay dishes (Corning 431111). Transformation efficiency was verified via serial dilution plates, and individual clones were validated through Sanger or whole-plasmid sequencing.

#### Functional cassette insertion and validation

A secondary cloning step was performed to insert the reporter cassette (minimal promoter-Cre-fluorophore-WPRE) into the barcoded library backbone. The cassette was PCR-amplified and inserted via restriction/ligation. To prevent backbone self-ligation, the library was treated with Shrimp Alkaline Phosphatase (rSAP) (NEB M0371S).

The final library was expanded via electroporation as described above. Library diversity and barcode distribution were confirmed by Next-Generation Sequencing (Illumina MiSeq, 2x250bp) of a 150–300 bp amplicon covering the barcode region (see Figure S6A) with an approach previously described.^64^ Validated pooled libraries were then processed for AAV packaging.

### AAV preparation

The pooled CRE library or individual AAV constructs were packaged into AAV-PHP.S at the Boston Children’s Hospital Viral Core or with PackGene Biotech, Inc. The titers (2–1000 × 10^12^ genome copies/mL) were determined by qPCR and adjusted to order requirements. The purity of the viruses was assessed via SDS-PAGE and the viruses were validated to be endotoxin-free via the Limulus Amebocyte Lysate (LAL) test.

### I.C.V. injections

For the intracerebroventricular (ICV) injection procedures in mice, AAV viruses at a dose of 1-4 × 10^10–11^ gc/ml along with Trypan Blue at a total injection volume of about 3ul were prepared. Neonatal pups at postnatal day 0-4 (P0-4) were cryo-anesthetized for 1-2 minutes and injected with the viruses to cerebral lateral ventricles using a Hamilton Gastight syringe (Hamilton 7653-01) with 32G needle. At least 4 weeks after AAV administration, DRG tissues were collected.

### RNAscope in situ hybridization

The mouse DRG dissections were performed as previously reported^65,66^ with minor modifications. In brief, mice were anesthetized with isoflurane for about 5 minutes and decapitated. The skin and muscles were removed from the dorsal aspect of the mice to expose the spinal cord (SC), then the vertebral columns of the SC were exposed and placed on a tray of ice. DRG were exposed after ½ vertebral columns were removed and the surrounding tissues were cut away. Finally, the collected DRGs were transferred to 4% PFA for a 24h incubation at 4°C, followed by a 24h incubation in 30% sucrose dissolved in PBS at 4°C. For sample preparation, DRG were stored at -80°C in Tissue-Tek O.C.T. compound (Electron Microscopy Sciences 62550-01) until they were cryosectioned. DRG were sectioned at 14 µm thickness, and mRNAs were detected by RNAscope (Advanced Cell Diagnostics, RNAscope® Fluorescent Multiplex Reagent Kit v1) using the manufacturer’s protocol. The following probes were used: *eGFP* (Cat# 400281), *Tac1* (Cat# 410351-C2), *Mrgprd* (417921-C2), *Rbfox3* (Cat# 313311-C3).

#### EGFP, Sstr2, Calca, Tubb3 Images

The mouse DRG dissections were performed as previously reported^67^ with minor modifications. In brief, mice were anesthetized with isoflurane for about 5min before cervical dislocation and decapitation. Fur was doused with 70% EtOH to restrict contamination. Pelt, skin, muscles, and viscera surrounding the spinal column were removed, and the spinal column was isolated, bisected, and placed in a dish with HBSS (Gibco 14175-095) on a tray of ice. Lumbar DRG were dissected and placed into cryomolds that were subsequently filled with Tissue-Tek O.C.T. compound (Electron Microscopy Sciences 62550-01) and allowed to solidify on dry ice for 5 minutes. DRG were stored at -80°C prior to cryosectioning. Lumbar DRGs were cryosectioned at 16 µm thickness and adhered to SuperFrost slides (Fisherbrand 12-550-15) and frozen at -80°C for storage or immediately processed. mRNAs were detected by RNAscope (Bio-Techne Advanced Cell Diagnostics, RNAscope® Fluorescent Multiplex Reagent Kit v2) using the manufacturer’s protocol. The following probes were used: *eGFP* (Cat# 400281-C1), *Sstr2* (Cat# 437681-C2), *Calca* (Cat# 557031-C3), and *Tubb3* (Cat# 423391-C4).

### Imaging and Quantification

Following FISH, sections were imaged with a Zeiss LSM 710 confocal microscope using the 20x objective for cell quantification and the 40x oil objective for showing cell markers at higher resolution. Cell quantifications were performed manually by ImageJ (Fiji, 2.3.0) using regions of interest (ROIs) to define the quantified area. Cell counts were confirmed by labeling with nuclei marker DAPI and pan-neuronal marker *Rbfox3*. AAV+ neurons were counted as cells with more than five *GFP*+ punctate signals per ROI. *Tac1*+ or *Mrgprd*+ cells were counted as cells with more than five *Tac1*+ or *Mrgprd*+ punctate signals, respectively, per ROI. The number of neurons that were either *Tac1*+*GFP*+ or *Mrgprd*+*GFP*+ were summed and then divided by the total number of *GFP*+ cells to obtain either %*Tac1*+*GFP*+ / *GFP*+ cells or %*Mrgprd*+*GFP*+ / *GFP*+ cells for each image per replicate. The values for either %*Tac1*+*GFP*+ / *GFP*+ cells or %*Mrgprd*+*GFP*+ / *GFP*+ cells were averaged across 2-5 images from a given animal and reported as a replicate. Replicates were obtained from 3-6 different individual mice.

#### EGFP, Sstr2, Calca, Tubb3 Images

Following FISH, sections were imaged with Andor DragonFly 600 Spinning Disk confocal microscope using the water immersion 25x objective. Cell quantifications were performed manually by ImageJ (Fiji, 2.17.0) using regions of interest (ROIs) to define the quantified area. Cell counts were confirmed by labeling with nuclei marker DAPI and pan-neuronal marker *Tubb3*. AAV+ neurons were counted as cells with more than five *EGFP*+ punctate signals per ROI. *Calca*+ or *Sstr2*+ cells were counted as cells with more than five *Calca*+ or *Sstr2*+ punctate signals, respectively, per ROI. The number of neurons that were either *EGFP*+*Calca*+*Sstr2*+ or *Calca*+*Sstr2*+ were summed and then divided by the total number of *EGFP*+ or *Calca*+ cells respectively, to obtain %*EGFP*+*Calca*+*Sstr2*+ / *EGFP*+ cells or %*Calca*+*Sstr2*+ / *Calca*+ cells for each image per replicate. The values for either %*EGFP*+*Calca*+*Sstr2*+ / *EGFP*+ cells or %*Calca*+*Sstr2*+ / *Calca*+ cells were averaged across 3-6 images from a given animal and reported as a replicate. Replicates were obtained from 3 different individual mice.

### DRG cell dissociation from enhancer AAV injected animals

Adult C57BL/6 mice were euthanized according to institutional guidelines. Spinal columns were dissected, bisected, and DRGs were carefully extracted from all axial levels in ice-cold Hank’s Balanced Salt Solution (HBSS; ThermoFisher Scientific, 14175103). Peripheral and central nerve roots were removed to isolate sensory ganglia, which were collected in Dulbecco’s Modified Eagle Medium (DMEM; ATCC, 30-2002) supplemented with 1% penicillin-streptomycin (P/S; Thermo Fisher, 15140122) on ice.

Ganglia were enzymatically dissociated using a Collagenase/Type I Trypsin/DNase I mixture (Sigma C9891, T8003; ThermoFisher Scientific 18047019) in DMEM at 37°C. Incubation times were adjusted based on age: 30 min for 4–6-week-old animals and 45 min for animals ≥8 weeks. Digestion was quenched with Neurobasal (NB) medium supplemented with 5% heat-inactivated fetal bovine serum (HI-FBS; Gibco, 16140063), 1% P/S, and 1X B27 supplement (Gibco, 17504044). Ganglia were triturated sequentially with fire-polished glass pipettes of decreasing bore size until a single-cell suspension was achieved, followed by filtration through a 70 µm nylon mesh (Pluristrainer, 43-10070-60). Cells were layered onto a 15% bovine serum albumin (BSA; Sigma, A9205) gradient and centrifuged at 130 × g for 10 min to remove debris. The pelleted cells were gently resuspended in NB medium, ensuring a final BSA concentration ≤2% to maintain compatibility with downstream single-cell applications (10x Genomics). Cell viability was assessed by trypan blue exclusion and hemocytometer counting. Preparations with >80% viable cells and minimal debris were used for downstream applications, including single-cell RNA sequencing.

### scRNA-seq of enhancer AAV libraries

Cell suspensions were sequenced using 10x Genomics assays, resuspended, and loaded into the 10x Chromium device for snRNA-seq (10x Genomics v3.1).

#### Targeted cDNA enrichment

Custom enrichment of targeted transcripts was performed using cDNA generated from the 10x Genomics Chromium Next GEM Single Cell 3’ RNA v3.1 workflow. For each enrichment reaction, 2 ng of cDNA template (obtained post-cDNA amplification) was amplified using Q5 High-Fidelity 2X Master Mix (New England Biolabs M0492S). The reaction was performed in a 25μL volume with a final concentration of 500 nM for each of the following primers:

● Forward (Partial Read 1): 5’-CTACACGACGCTCTTCCGATCT-3’
● Reverse: 5’-TGTGCACTGTGTTTGCTGACGC-3’

The PCR amplification was conducted with an initial denaturation at 98°C for 30 seconds, followed by 15 cycles of denaturation at 98°C for 10 seconds, annealing at 69°C for 20 seconds, and extension at 72°C for 30 seconds. A final extension was performed at 72°C for 2 minutes.

Post-amplification, the enriched product was purified using the QIAquick PCR Purification Kit (Qiagen) and eluted in a volume of 22μL EB. Product integrity and size distribution were assessed via Agilent TapeStation. The purified, enriched product was integrated back into the standard 10x Genomics workflow starting at Step 3.1 (Gene Expression Library Construction). To account for the specific nature of the enriched fragments, the enzymatic fragmentation time was reduced from 5 minutes to 2 minutes. All subsequent steps, including adapter ligation and index PCR, were completed according to the Chromium Next GEM Single Cell 3’ Reagent Kits v3.1 (Dual Index) User Guide.

All other libraries were prepared for scRNA-seq according to the manufacturer’s protocol. Libraries were sequenced on an Illumina NextSeq 2000 or shipped to GENEWIZ (Azenta Life Sciences) for sequencing on the Illumina NovaSeq X Plus platform to an approximate read depth of ∼20,000 reads per cell. Our scRNA-seq analysis pipeline followed a protocol similar to our previous studies;^6,61^ the key steps for the single virus and AAV CRE libraries are detailed below.

#### Single-cell RNA-seq processing of CRE libraries

All single-cell libraries (single-virus CREs and AAV CRE libraries) were aligned to the mm10 genome (modified to include the viral vector backbone sequence) using 10x Genomics CellRanger v9. Ambient RNA was removed using CellBender v0.3.0 (single-virus libraries: v0.3.0), using default parameters for both CellRanger and CellBender. For the AAV CRE library, viral barcode detection was performed from CellRanger BAM outputs. Viral barcoded reads were extracted using custom Python scripts, and annotated by matching read sequences to the known CRE barcode list (allowing a single base-pair mismatch).

For both the single-virus and AAV CRE libraries, ambient RNA-corrected filtered gene count matrices were processed in Seurat v4. Low quality cells were excluded (detected genes > 500; mitochondrial reads < 20%). Data were then split by sequencing library and integrated across runs using Seurat’s standard workflow as described in Bhuiyan et al., 2024. Clustering was performed at resolution = 1.0, and clusters were annotated using established marker genes for neuronal (*Snap25*, *Rbfox3*) and non-neuronal populations (e.g., *Sparc*, *Ptprc*, *Ptgfra*).

For single-virus libraries, a second round of integration was performed. The neuronal population was subsetted and reintegrated using the same pipeline with a stricter QC threshold (detected genes > 1,000). Final neuronal identities were assigned by anchoring to the Bhuiyan et al., 2024^6^ mouse DRG reference atlas using Seurat v4 label transfer; predicted labels were used as final annotations.

For the AAV CRE library, transcriptomic processing, filtering, integration, clustering, and marker-based annotation were performed as above, followed by Seurat v4 label transfer to the Bhuiyan et al., 2024^6^ DRG mouse neuron-only reference atlas. Predicted labels were used as final annotations, except for clusters enriched for *Sparc* and *Mpz*, which were classified as non-neuronal.

To quantify CRE activity across functional neuronal classes, a cell class specificity ratio was computed for each CRE as: number of CRE barcode–positive cells in a given DRG cell class divided by number of barcode–positive cells across all other cell classes. DRG cell classes were defined by aggregating transcriptomic clusters into: C-PEP (Calca+Sstr2, Calca+Dcn, Calca+Oprk1, Calca+Adra2a), C-NP.Mrgpra3 (Mprgra3+Trpv1, Mrgpra3+Mrgprb4), C-NP.Mrgprd, C-NP.SST, A-LTMR (Ntrk3^high^+Ntrk2, Ntrk3^high^+S100a16, Ntrk3^low^+Ntrk2), A-PEP (Calca+Smr2, Calca+Bmpr1b), A-Propr, C-LTMR.Th, C-Thermo (Trpm8 and Rxfp1), and non-neurons.

To define the “on-target” cell type for each CRE, candidate on-target sets (e.g., C-NP.Sst, C-Thermo.Trpm8) were assigned based on accessible peak evidence from UCSC Genome Browser tracks (downsampled to 270 cells per cell type). A DA score was then calculated by comparing chromatin accessibility in the proposed on-target versus off-target cell types. DA was computed using a logistic regression test (test.use = “LR”) with peak counts as a latent variable (latent.vars = “nCount_MACS”). On-target assignments were designated “highly likely” (Table S16) when the DA value (log2FC on-target vs off-target) was positive.

### PAIN-net development

To decode the regulatory logic determining cell-type specificity, we developed PAIN-net, a deep convolutional neural network (CNN) framework that predicts chromatin accessibility and differential expression directly from DNA sequence. We chose this architecture because CNNs can learn non-linear relationships, both between input sequences and between input sequences and the predicted variable(s), and CNNs do not require explicit featurization. Consequently, previous studies have shown that CNNs sufficiently infer CRE function from sequences.^43,69^

#### Model architecture

PAIN-net utilizes a 1D Residual Network (ResNet) architecture to process 800-bp input sequences. The ResNet architecture was chosen for its ability to learn complex, high-order motif combinations while mitigating gradient degradation in deep models.^70,71^ The network is composed of an initial convolutional layer followed by five stacked residual blocks. To integrate sequence information across broader genomic contexts, we employed dilated convolutions within these blocks with exponentially increasing dilation rates. This design expands the receptive field to capture long-range interactions while preserving single-nucleotide resolution, as demonstrated in previous genomic models like Basenji.^71^ The feature maps are aggregated via global average pooling and fed into a shared dense layer. This layer bifurcates into two task-specific heads: one predicting differential expression (log2-fold change) and the other predicting chromatin accessibility (ATAC-seq signal from bigwig files) across cell types.

#### Data preprocessing and encoding

To identify subtype-specific regulatory elements, we performed differential accessibility analysis on single-cell ATAC-seq data using the FindAllMarkers function in Signac.^72^ We compared each neuronal subtype against all other subtypes to identify differentially accessible sequences. The analysis was conducted using default parameters with specific adjustments to account for technical covariates and capture a broad range of fold-changes: the logistic regression test (test.use = “LR”) was employed with peak counts as a latent variable (latent.vars = “nCount_MACS”) and the log2-fold change threshold set to zero (log2fc.threshold = 0). The resulting log2-fold change (log_2_FC) values served as the primary regression targets for the differential expression head of the model. Additionally, to capture absolute chromatin accessibility levels, each sequence was annotated with raw fragment number per base.

Genomic sequences were centered on the identified peaks and fixed to a length of 800 base pairs (bp). DNA sequences were one-hot encoded into an 800 x 4 binary matrix. To improve model invariance to strand orientation and increase the effective training set size, data augmentation was performed by including the reverse complement of all training sequences. Target labels (log_2_FC and ATAC signal) were standardized to zero mean and unit variance per cell type to ensure gradient stability during training.

#### Model training and optimization

The model was trained using a multi-objective loss function. We minimized a hybrid loss combining Mean Squared Error (MSE) and Pearson correlation (r) for the expression head (L = MSE - r). This ensured that the model prioritized both the magnitude of fold-changes and the relative expression ranking across cell types. A standard MSE loss was applied to the chromatin accessibility head as an auxiliary task to regularize the shared representation.

To mitigate the sparsity of rare neuronal subtypes, we implemented a dynamic weighting strategy. Loss contributions were balanced by the inverse frequency of available data per cell type. Furthermore, we applied targeted boosting weights (2x) to biologically critical but underrepresented clusters (i.e peptidergic subtypes) while dampening dominant signals to prevent overfitting to majority classes (0.5x; i.e Atf3). Training was performed using the Adam optimizer with a learning rate of 10^-3^. We utilized early stopping based on the mean Pearson correlation of the validation set to prevent overfitting.

#### In silico mutagenesis

We performed *in silico* saturation mutagenesis (ISM) on CRE1-115 to quantify position- and nucleotide-specific contributions to predicted enhancer activity across all modeled cell types. For each CRE, we extracted an 800-bp sequence window centered on the peak summit by taking 400 bp of flanking sequence upstream and 400 bp downstream of the peak center, and one-hot encoded each sequence into an 800×4 matrix (A/C/G/T). The PAIN-net model was used to generate predictions, and target-specific output scalers (loaded from a pickled dictionary) were applied to inverse-transform model outputs back to the original scale (reported as log_2_FC). For each sequence, a baseline prediction was computed for the wild-type input, followed by generation of all single-nucleotide variants at every position (3 substitutions per valid base; 800 positions; 18 cell types). For each cell type, an ISM matrix of shape 4×800 was constructed where each entry represents the change in predicted activity (Δlog2FC) for substituting a given base at a given position relative to the wild-type prediction. Per-CRE outputs were saved into a hierarchical directory structure (CRE → target), including (i) fixed-scale heatmaps of the ISM matrix (−1 to 1), (ii) per-mutation scatterplots of Δlog2FC versus position colored by mutant base (−1 to 1), and (iii) attribution-style sequence logos derived by assigning the wild-type base at each position the negative mean effect of its alternative substitutions (−1 to 1). In addition, for each target we computed an “importance profile” defined as the maximum absolute Δlog2FC across the four bases at each position, and aggregated these profiles across all CREs to produce global summary heatmaps (0 to 1) showing position-wise importance per sequence for each target.

### Electrophysiological data acquisition and analysis

#### Primary sensory neuron culture

35-mm culture dishes were sequentially coated with Poly-D-Lysine (1–2 h, 37°C) and laminin (2.5 µg/mL), then air-dried prior to plating. Dorsal root ganglia (DRGs; ∼35 per mouse) were harvested from wild-type mice (≥postnatal day 10) into ice-cold DMEM/F12. Tissue was enzymatically digested in HBSS containing Collagenase A (2 mg/mL) and Dispase (2 mg/mL) for 60 min at 37°C, followed by mechanical trituration to a single-cell suspension. To eliminate myelin and debris, the suspension was centrifuged (200 rcf, 12 min) through a 30% BSA density gradient. The resulting neuronal pellet was resuspended in complete Neurobasal medium (B27, Glutamax, Pen/Strep) and seeded at ∼100,000 cells/dish. For experiments requiring K_ir_2.1 expression, neurons were transduced *in vitro* immediately following dissociation. Control neurons were not transduced. AAV vectors (CAG-K_ir_2.1-mCherry, or CRE1-K_ir_2.1-mCherry) were applied at a titer of 1 x 10^13^ vg/mL. Specifically, 1–2 microliters of viral stock (1–2 x 10^10^ vg) was added to 5 x 10^5^ cells in a 200 μL volume. Cultures were incubated overnight, and electrophysiological recordings were performed 24–48 hours post-dissociation to ensure robust ion channel expression.

#### Recording configurations and quality control

Whole-cell patch-clamp recordings were performed 24–48 h post-dissociation using an EPC 10 amplifier (HEKA) at room temperature. Small-diameter nociceptive neurons (<25μm, capacitance<15pF) were targeted. Recording electrodes (5–6 MΩ) were filled with a K-gluconate-based internal solution (see below). Following giga-ohm seal formation, series resistance (10–15 MΩ) was compensated by ≥80%. Data were sampled at 20 kHz and low-pass filtered at 5 kHz.

#### Electrophysiological *solutions*

The standard extracellular bath solution contained (in mM): 155 NaCl, 3.5 KCl, 1.5 CaCl_2_, 1 MgCl_2_, 10 HEPES, and 10 glucose (pH 7.4 adjusted with NaOH). The intracellular pipette solution consisted of (in mM): 140 K-gluconate, 13.5 NaCl, 1.8 MgCl_2_, 0.09 EGTA, 9 HEPES, 14 creatine phosphate (Tris salt), 4 MgATP, and 0.3 Tris-GTP (pH 7.2 adjusted with KOH).^73^

### Stimulation protocols and data analysis

The resting membrane potential (RMP) was measured in current-clamp mode immediately after achieving the whole-cell configuration without current injection. Cell capacitance was determined by the amplifier’s automated compensation circuit. In voltage-clamp mode, total ionic currents were elicited from a holding potential of −60 mV using 500-ms voltage steps ranging from −120 to -40 mV in 10 mV increments, with a 1s interpulse interval. Neuronal excitability was assessed in current-clamp mode using 2-s depolarizing steps (10–50 pA increments). Rheobase was defined as the minimum current required to elicit an action potential.

While CAG-Kir2.1-mCherry neurons were identified by robust fluorescence, CRE1-driven expression was often indistinguishable from autofluorescence. To objectively identify successfully transduced cells in the CRE1 group, we utilized a functional inclusion criterion based on neuronal excitability. Based on an initial characterization of control neurons, which exhibited a consistent rheobase (Mean: 55.5 pA), we established a threshold for “High Rheobase” cells. Cells in the experimental groups (CRE1-K_ir_2.1-mCherry) were categorized as CRE1-K_ir_2.1-mCherry cells in Figure 6 if they exhibited a rheobase >200 pA (Figure S8A). This stringent threshold (representing a >3.5-fold increase over the control mean) served as a physiological proxy for successful K_ir_2.1 expression, as the increased membrane leak conductance provided by K_ir_2.1 is known to shunt depolarizing currents and increase the threshold for action potential firing.^74^

Current densities (pA/pF) were calculated by normalizing peak or steady-state currents to cell capacitance. Data were analyzed using custom MATLAB scripts and OriginPro, with statistical comparisons (unpaired t-tests) performed in GraphPad Prism 9. Results are presented as mean± SEM.

### Human iPS-derived nociceptors and cardiomyocytes cell culture assay

The human iPS cell-derived nociceptors and cardiomyocytes cell culture were performed as described here: iPS cell-derived nociceptors were plated at 5,000/well density and co-cultured with 2,000 glial cells in 384-well plate, iPS cell-derived cardiomyocytes were plated at 10,000/well without co-culture. Human iPS cells were infected by directly adding concentrated AAV viral particles with multiple titers: 1x10^9^, 1x10^10^, 1x10^11^, 1x10^12^ to a well plate, two well plate replicates per group. Cells were re-fed every two days with maintenance medium (Nociceptors: DMEM/F-12 Glutamax (ThermoFisher Scientific 10565018), N-2 Supplement (ThermoFisher Scientific A1370701), B-27 Supplement (ThermoFisher Scientific A3353501), BDNF (Bio-Techne R&D Systems 248-BDB-050/CF), GDNF (Bio-Techne R&D Systems 212-GD-050/CF), ß-NGF (Bio-Techne R&D Systems 256-GF-100/CF), NT-3 (Bio-Techne R&D Systems 267-N3-025/CF); Cardiomyocytes: RPMI 1640 medium (ThermoFisher Scientific 11835055), 1x B27 supplement with insulin (ThermoFisher Scientific 17504-044)). After culturing for two weeks, cells were fixed with 4% PFA for 0.5h and rinsed with PBS three times. Following that, the images were acquired with a Zeiss LSM 710 confocal microscope using the 10x objective and 40x objective for zoom-in images.

## DATA AND CODE AVAILABILITY

All custom code used to integrate data is available on Git (https://github.com/Renthal-Lab/painseq-multiome). Raw sequencing data and final objects will be deposited into GEO.

## Figure Legends

**Figure S1:**
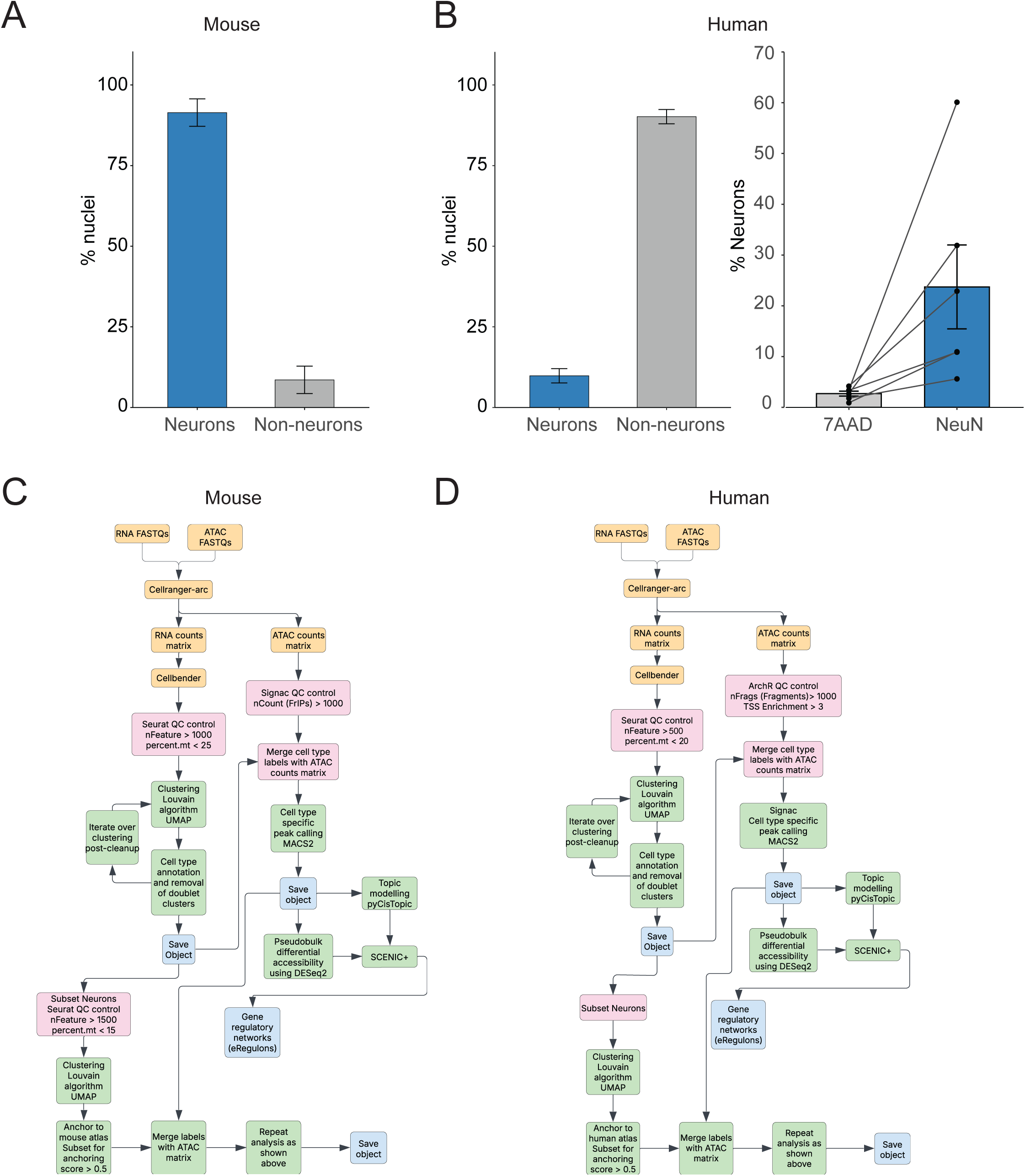
Analysis pipeline and RNA-level quality control metrics for mouse and human DRG cell types. **A:** Barplot displays the fraction of neurons and non-neurons obtained from the multi-omic sequencing of *Sun1-GFP+/-;vGlut2-Cre+/-* sorted mouse DRG nuclei. Error bars represent standard error across libraries. **B:** (*Left*) Barplot displays fraction of neurons and non-neurons obtained from multi-omic sequencing of human DRG nuclei. Error bars represent the standard error across samples. (*Right*) Barplot displays the fraction of neuronal enrichment in each sample that was sorted either for a high 7AAD signal or both high 7AAD and NeuN signals. Lines connect measurements from the same human DRG samples (dots) using each sorting strategy. **C:** Flowchart displays the bioinformatic pipeline detailing how snMultiome data was processed for mouse DRG. See Methods for details. Abbreviations: ATAC (assay for transposase-accessible chromatin); QC (quality control); nFeature (number of detected genes per cell); percent.mt (percentage of mitochondrial reads); FrIP (fraction of reads in peaks); nFrags (number of ATAC fragments per cell); TSS (transcription start site); UMAP (uniform manifold approximation and projection); eRegulon (enhancer-driven regulatory module) **D:** Flowchart displays the bioinformatic pipeline detailing how snMultiome data was processed for human DRG. See Methods for details.

**Figure S2:**
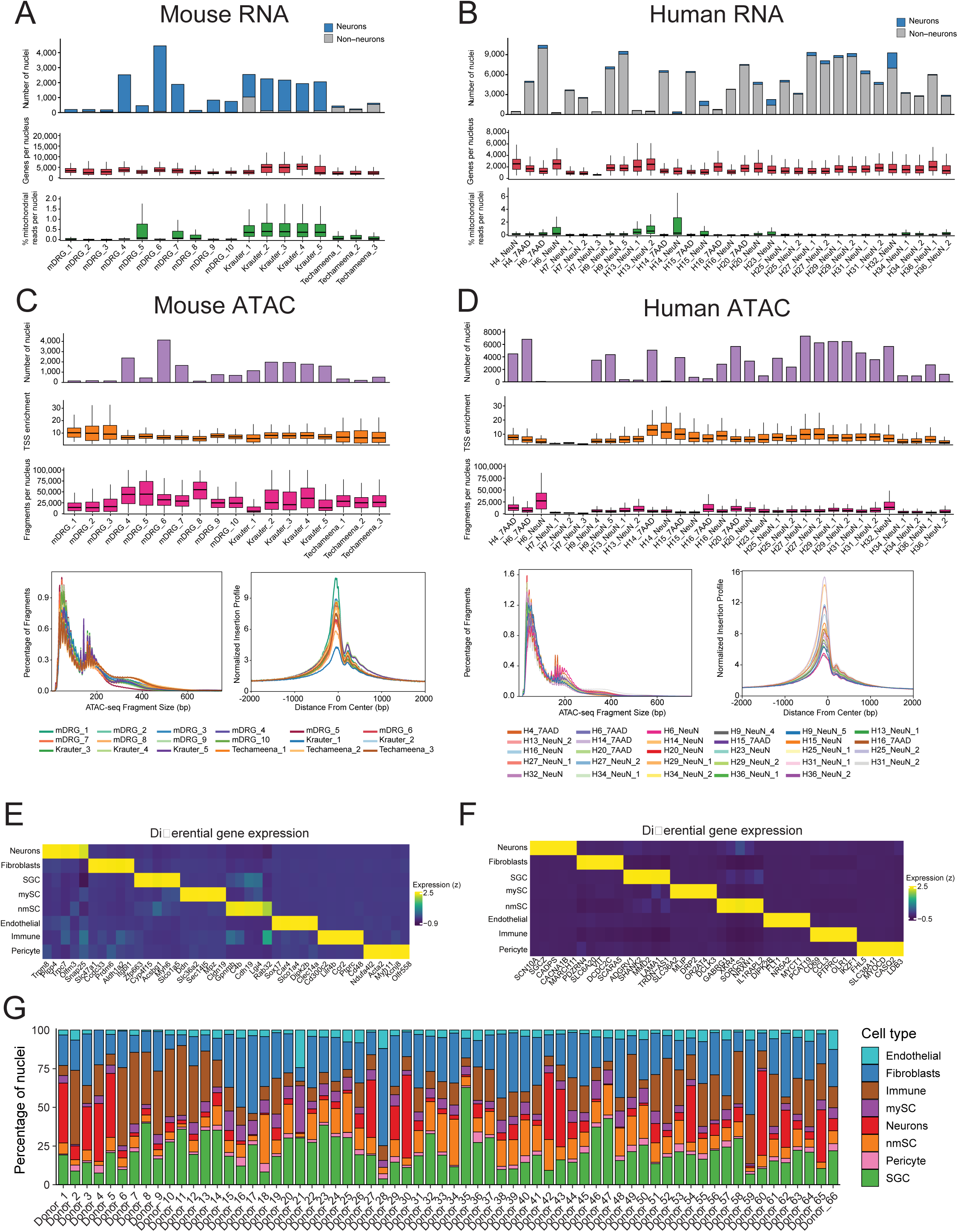
snMultiome level (RNA+ATAC) quality control metrics for mouse and human DRG cell types. **A**: Plots display RNA-level snMultiome metrics for mouse DRG nuclei that passed quality controls and were included in downstream analyses. (*Top*) Stacked bar plots display the number of neuronal (blue) and non-neuronal (gray) nuclei that passed quality controls by sample. (*Middle*) Box plots display the number of unique genes detected per nucleus in each sample. (*Bottom*) Box plot displays the percentage of mitochondrial reads detected per nucleus in each sample. The median value is indicated by the horizontal black line inside the box. Boundaries of the boxes mark quartiles, and whiskers represent 1.5 times the interquartile range (Q1–Q3). Krauter and Techameena refer to previous publications, Krauter et al., 2025^20^ and Techameena et al., 2024,^19^ respectively. **B:** Plots display RNA-level snMultiome metrics for human DRG nuclei that passed quality controls and were included in downstream analyses. (*Top*) Stacked bar plots display the number of neuronal (blue) and non-neuronal (gray) nuclei that passed quality controls by sample. (*Middle*) Box plots display the number of unique genes detected per nucleus in each sample. (*Bottom*) Box plot displays the percentage of mitochondrial reads detected per nucleus in each sample. The median value is indicated by a black line inside the box. Boundaries of the boxes mark quartiles, and whiskers represent 1.5 times the interquartile range (Q1–Q3). **C:** Plots display ATAC-level snMultiome metrics for mouse DRG nuclei that passed quality controls and were included in downstream analyses. (*Top*) Bar plot displays the number of nuclei that passed quality controls, colored by sample. (*Middle*) Box plots display TSS enrichment scores per nuclei for each sample. (*Bottom*) Box plots display unique fragments detected per nuclei in each sample. The median value is indicated by a black line inside the box. Boundaries of the boxes mark quartiles, and whiskers represent 1.5 times the interquartile range (Q1–Q3). (*Bottom, Left*) Histogram displays the fragment size distribution of ATAC-seq fragments for each sample, with the y-axis representing the percentage of fragments per sample. Lines are colored by sample. (*Bottom, Right*) Tn5 insertion profile displays normalized insertion signal around transcription start sites for each sample (x-axis shows distance from TSS center in bp; y-axis shows normalized insertion profile). **D:** Plots display ATAC-level snMultiome metrics for human DRG nuclei that passed quality controls and were included in downstream analyses. (*Top*) Bar plot displays the number of nuclei that passed quality controls, colored by sample. (*Middle*) Box plots display the distribution of TSS enrichment scores per cell for each sample. (*Bottom*) Box plots display the distribution of unique fragments detected per cell for each sample. The median value is indicated by a black line inside the box. Boundaries of the boxes mark quartiles, and whiskers represent 1.5 times the interquartile range (Q1–Q3). (*Bottom, Left*) Histogram displays the fragment size distribution of ATAC-seq fragments for each sample, with the y-axis representing the percentage of fragments per sample. Lines are colored by sample. (*Bottom, Right*) Tn5 insertion profile displays the normalized insertion signal around transcription start sites for each sample (x-axis shows distance from TSS center in bp; y-axis shows normalized insertion profile). **E:** Heatmap displays z-scaled, log-normalized expression of marker genes (columns) for each cell type (rows) in mouse DRG. Marker genes are defined as differentially expressed in each cell type compared to all other (Log_2_FC > 0.5, adj. p-val < 0.05). **F:** Heatmap displays z-scaled, log-normalized expression of marker genes (columns) for each cell type (rows) in human DRG. Marker genes are defined as differentially expressed in each cell type compared to all others (Log_2_FC > 0.5, adj. p-val < 0.05). **G:** Stacked barplot displays the distribution of DRG cell types annotated in each donor.

**Figure S3:**
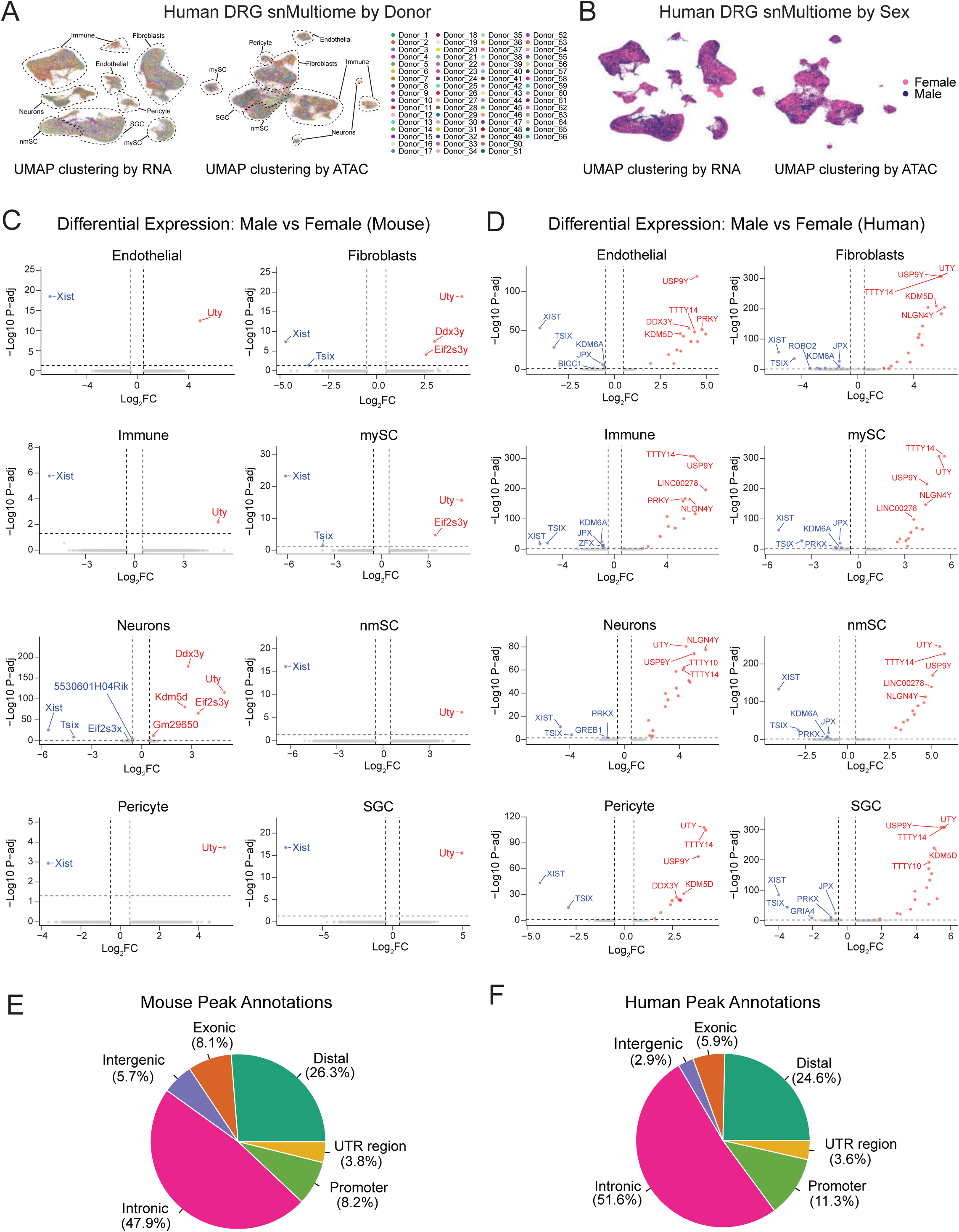
Donor-level integration and sex-specific differences in human DRG cell types. **A:** (*Left*) UMAP projection clustered by gene expression (same as in Figure 1C) displays 145,562 human DRG nuclei that passed RNA quality controls. Nuclei are colored by donor ID. DRG cell types are outlined (dashed outlines). (*Right*) UMAP projection clustered by chromatin accessibility displays 67,025 human DRG nuclei that passed ATAC quality controls. Nuclei are colored by donor ID. DRG cell types are outlined (dashed outlines) and annotated. **B**: (*Left*) UMAP projection clustered by gene expression (same as in Figure 1C) displays 145,562 human DRG nuclei that passed RNA quality controls. Nuclei are colored by sex. DRG cell types are as in A. (*Right*) UMAP projection clustered by chromatin accessibility displays 67,025 human DRG nuclei that passed ATAC quality controls. Nuclei are colored by sex. DRG cell types are as in A. **C:** Volcano plots display the log_2_FC (x-axis) and adj. p-values (y-axis) of genes differentially expressed in mouse males compared to females for each DRG cell type (log_2_FC > 0.5, adj. p-value < 0.05; threshold indicated by dotted black lines). **D**: Volcano plots display the log_2_FC (x-axis) and adj. p-values (y-axis) of genes differentially expressed in human males compared to females for each DRG cell type (log_2_FC > 0.5, adj. p-value < 0.05; threshold indicated by dotted black lines). **E:** Pie chart displays the fraction of MACS2-called peaks in mice DRG that are located in distinct genomic regions, as annotated by ChIPSeeker.^75^ Peaks within the promoter window were defined as those falling −1 kb to +100 bp relative to the annotated TSS. Peaks were annotated as Distal if they lay within ±200 kb of a TSS and are not promoters, or as Intergenic if they were ≥200 kb from the nearest TSS. **F:** Pie chart displays the fraction of MACS2-called peaks that are located in distinct genomic regions, as annotated by ChIPSeeker. Genomic regions defined as in E.

**Figure S4:**
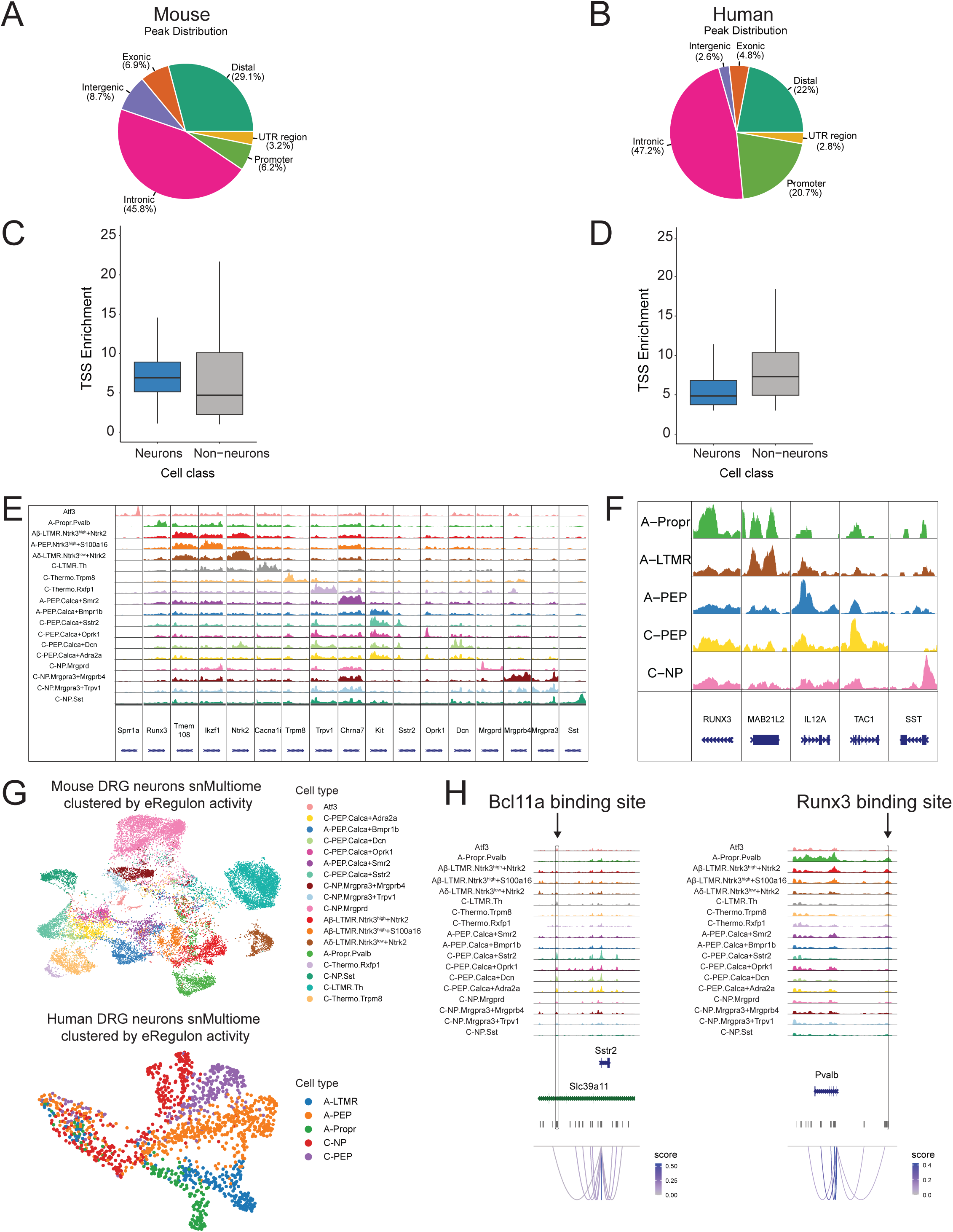
QC plots for snMultiome analysis of mouse and human DRG neuronal subtypes. **A**: Pie chart displays the fraction of MACS2-called peaks from mouse DRG neurons located in distinct genomic regions, as annotated by ChIPseeker. Promoter peaks are defined as those falling within −1 kb to +100 bp of the annotated TSS. Distal peaks are defined as regions within ±200 kb of a TSS but are not promoters, while Intergenic peaks are those ≥200 kb from the nearest TSS. **B**: Pie chart displays the fraction of MACS2-called peaks from human DRG neurons located in distinct genomic regions, as annotated by ChIPseeker. Genomic regions are defined as in A. **C:** Boxplot displays TSS enrichment scores per nucleus in mouse DRG neurons and non-neurons that passed all QC filters. TSS enrichment scores are defined as the fold change in read density at transcription start sites relative to the average background signal in the 100 bp distal flanking regions. **D:** Boxplot displays TSS enrichment scores per nucleus in human DRG neurons and non-neurons that passed all QC filters. TSS enrichment scores are defined as the fold change in read density at transcription start sites relative to the average background signal in the 100 bp distal flanking regions. **E:** Genomic tracks display chromatin accessibility at loci of canonical marker genes for each mouse DRG neuronal subtype. Accessibility at each locus (y-axis) is scaled to the max value across all cell types (rows). Gene bodies for each marker gene are displayed along the x-axis. **F:** Genomic tracks of chromatin accessibility at loci of marker genes for each human DRG neuronal subtype. Accessibility at each locus (y-axis) is scaled to the max value across all cell types (rows). Gene bodies for each marker gene are displayed along the x-axis. **G:** UMAP projection displays 18,238 mouse (*Top*) and 1,597 human (*Bottom*) nuclei clustered by their eRegulon scores (see Methods). Nuclei are colored by cell type annotations (as in Figure 2). **H:** Genomic tracks display chromatin accessibility of a candidate SCENIC+ CRE upstream of *Sstr2* in mice (left) and *PVALB* in humans (right). Boxed region indicates that the CREs are predicted binding sites for *BCL6* (mice) and *RUNX3* (humans). Loops show Pearson correlation (p-val. < 0.05) of enhancer accessibility and RNA expression of *BCL6* or *RUNX3*.

**Figure S5:**
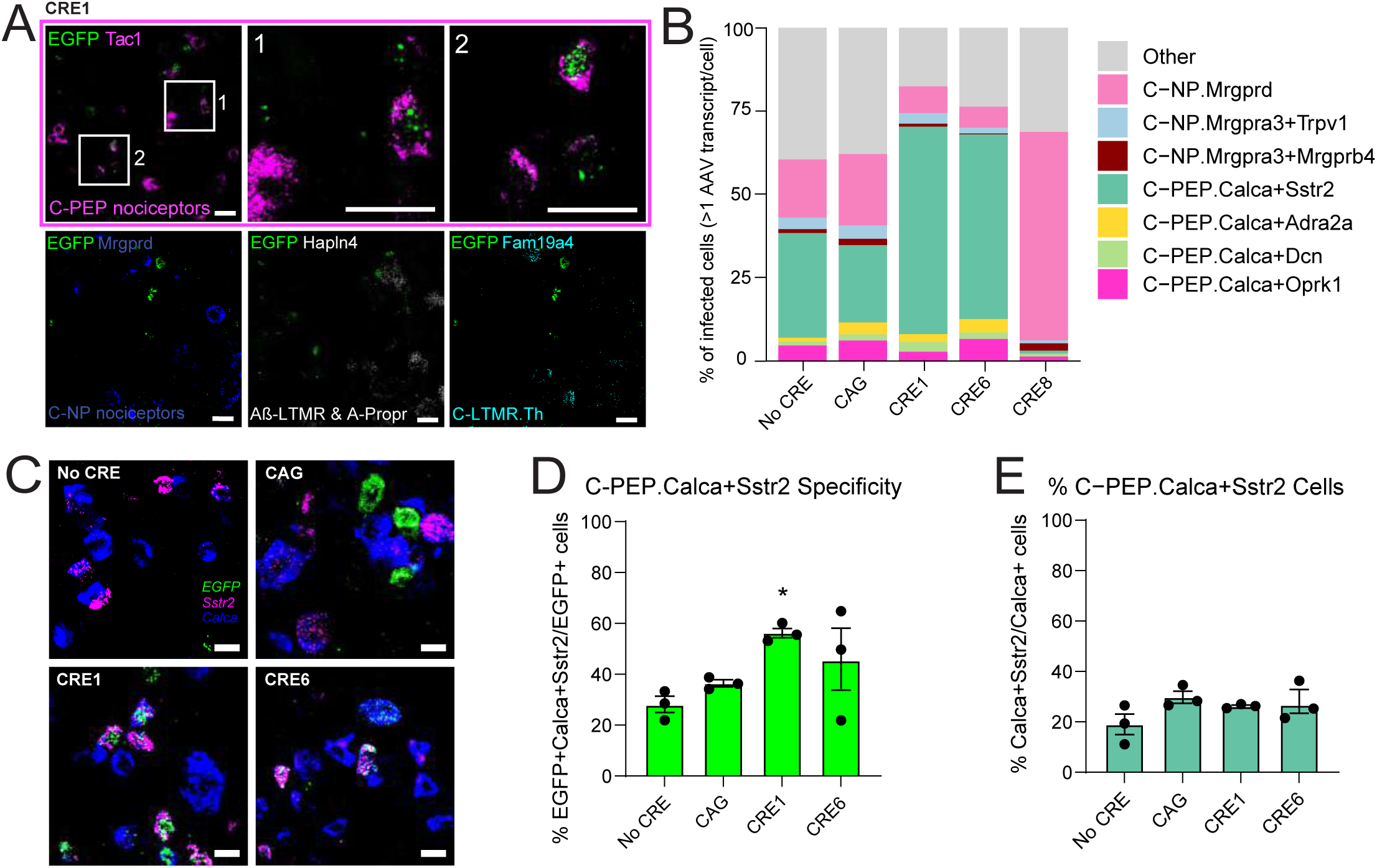
Screening candidate CREs for C-PEP.Calca+Sstr2 specificity. **A**: Representative images of mouse DRG neurons labeled by RNAscope *in situ* hybridization respectively for *Tac1* (magenta), *Mrgprd* (blue), *Hapln4* (white), *Fam19a4* (red), and viral *EGFP* (green) mRNA for CRE1, with insets illustrating colocalization of *EGFP* and *Tac1* for CRE1. Scale bar, 25 µm. **B:** Stacked bar plots of the fraction of each DRG neuronal subtype that express the respective CRE barcodes (>1 AAV transcript/cell). Colors correspond to C-NP or C-PEP subtypes; all other cell types are gray. **C:** Representative images of mouse DRG neurons infected with AAVs using NoCRE, CAG, CRE1, or CRE6 to drive *EGFP* expression. DRG neurons are labeled by RNAscope *in situ* hybridization for *Calca* (blue), *Sstr2* (magenta) and *EGFP* (green) mRNA. Scale bar, 25 µm. Quantification in D. **D:** Quantification of *Calca, Sstr2,* and *EGFP* mRNA colocalization per *EGFP+* cell for No CRE, CAG, CRE1, and CRE6 (n = 3-4 lumbar DRGs per mouse from 3 animals). One-way ANOVA, F(3,8) = 2.005, p = 0.0404, Dunnett’s multiple comparisons test for No CRE versus CRE1. Data are presented as mean ± SEM. **E:** Quantification of C-PEP.Calca+Sstr2 cells as a percentage of *Calca+* cells that are also *Sstr2+*. Data are presented as mean ± SEM.

**Figure S6:**
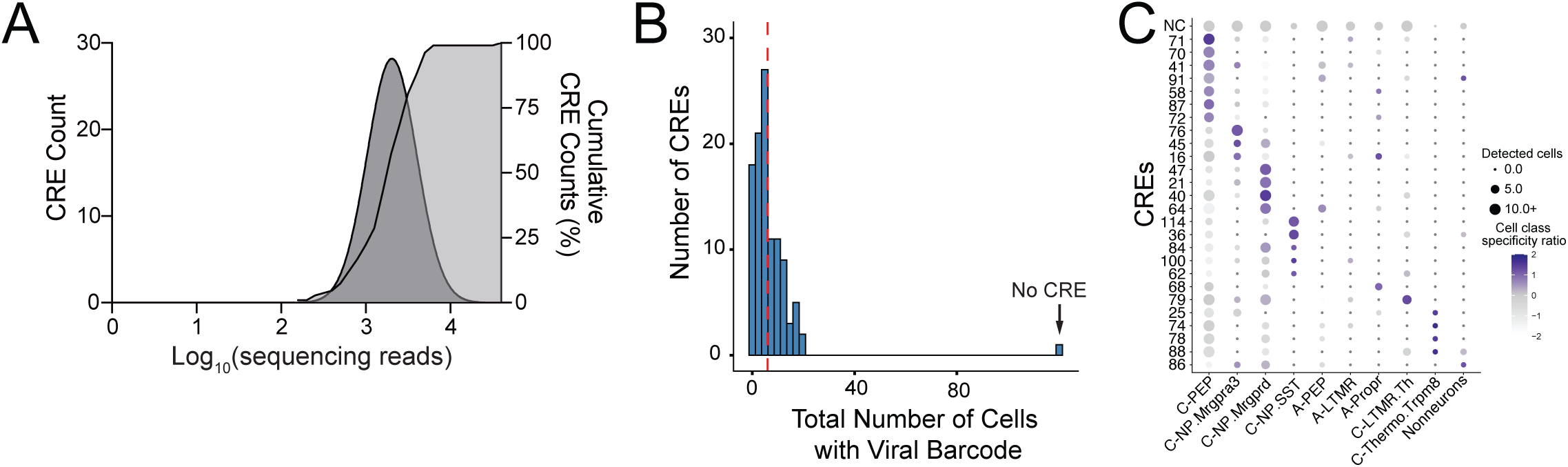
Characterization of CRE Library Complexity, Barcode Recovery, and Chromatin Accessibility. **A:** Distribution of CRE barcodes detected in the library AAV multiplexed CRE library as measured by next-generation DNA sequencing. Reads were binned by their associated CRE barcode (left y-axis) and their frequency distribution is plotted (dark gray). Cumulative distribution of detected CRE barcodes indicates coverage of all 106 prioritized CREs, the mini CMV enhancer, and no CRE negative controls (light gray, right y-axis) in the library. **B:** Histogram of number of DRG cells expressing viral barcodes from the multiplexed CRE library as measured by scRNA-seq (106 CREs + 2 controls). Only CREs detected in at least 6 cells were used (red line). No CRE control was over-represented in the library to ensure robust comparison for each CRE. **C:** Dotplot displays cell class specificity ratios across 10 distinct DRG cell classes. The cell class specificity ratio (x-axis) is defined as the number of CRE barcode-positive cells in a specific cell class divided by the number of barcode-positive cells in all other cells. Dot color represents cell class specificity ratio (Z-scaled), and dot size corresponds to the number of viral barcodes detected per cell type. Ratios are normalized to the negative control (NC: No CRE).

**Figure S7:**
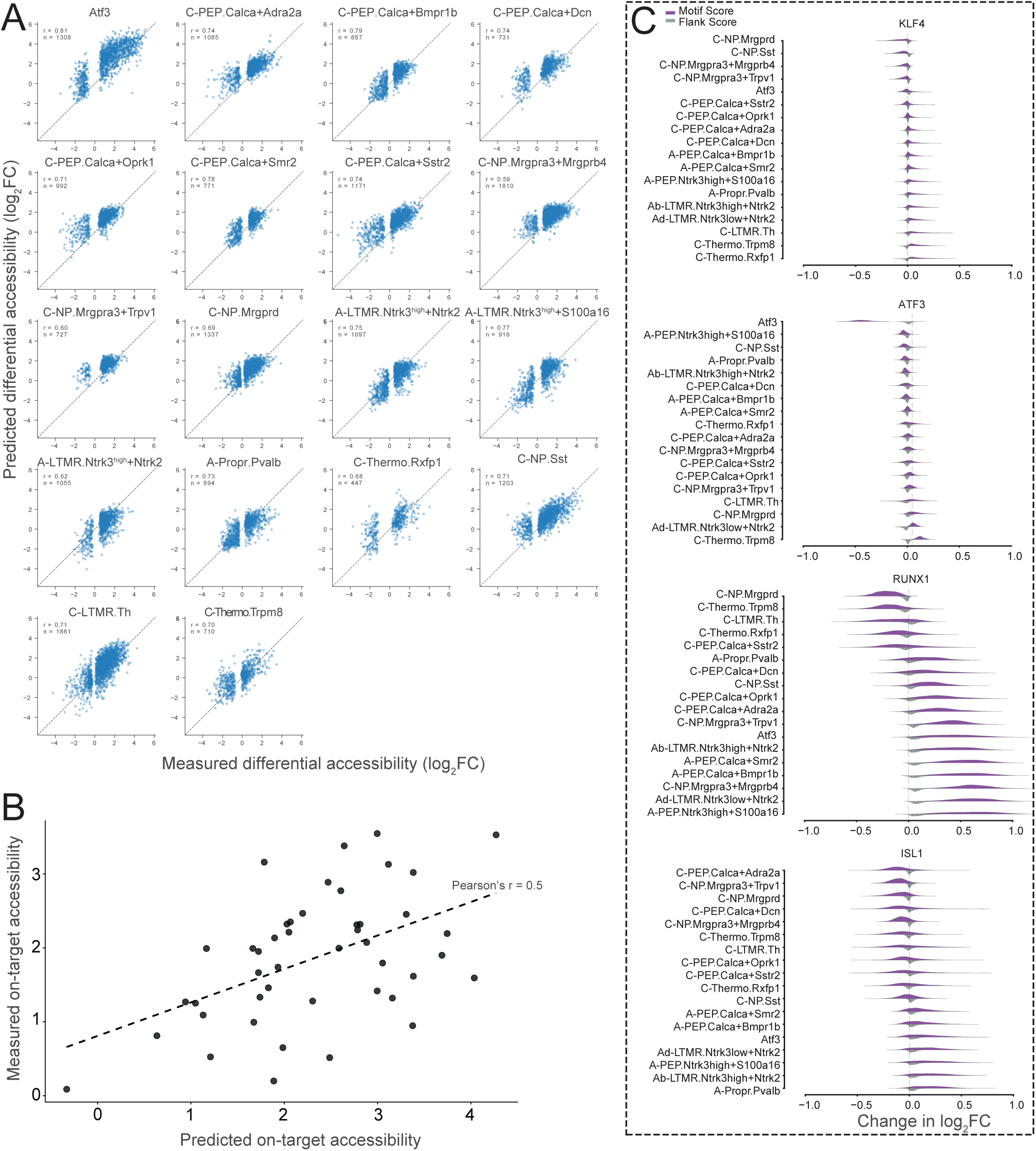
Constructing PAIN-net to aid in the prioritization of DRG CREs. **A:** Scatter plots compare experimentally measured (x-axis) versus predicted (y-axis) log_2_FC of each neuronal subtype compared to all others. Data points are all held-out validation sequences (15%). Each subplot reports the Pearson correlation (*r*) and sequence count (*n*). The dashed diagonal line (y=x) indicates perfect prediction accuracy. **B:** PAIN-net predicted on-target accessibility of candidate CRE sequences correlates with their measured on-target accessibility. On-target cell types are defined by the accessibility at each CRE locus in the multiplexed CRE library. CREs were designed to target multiple neuronal subtypes, so predicted and measured on-target accessibilities were summed across their on-target subtypes (Table S16). **C:** PAIN-net predicts bases that drive cell type-specific differential accessibility in motifs of cell-type-specific transcription factors. Split violin plots display the distributions of predicted mutation impacts (delta log_2_FC in each cell type compared to all other cell types) for the core consensus motif (purple) and the 10 bp flanking sequence (gray), aggregated across 1,000 randomized genomic backgrounds. Plots represent the aggregate results for *Atf3, Runx1, Jun,* and *Isl1* across DRG cell types (sorted by effect size). The X-axis displays the mean change in accessibility across the three possible mutations.

**Figure S8:**
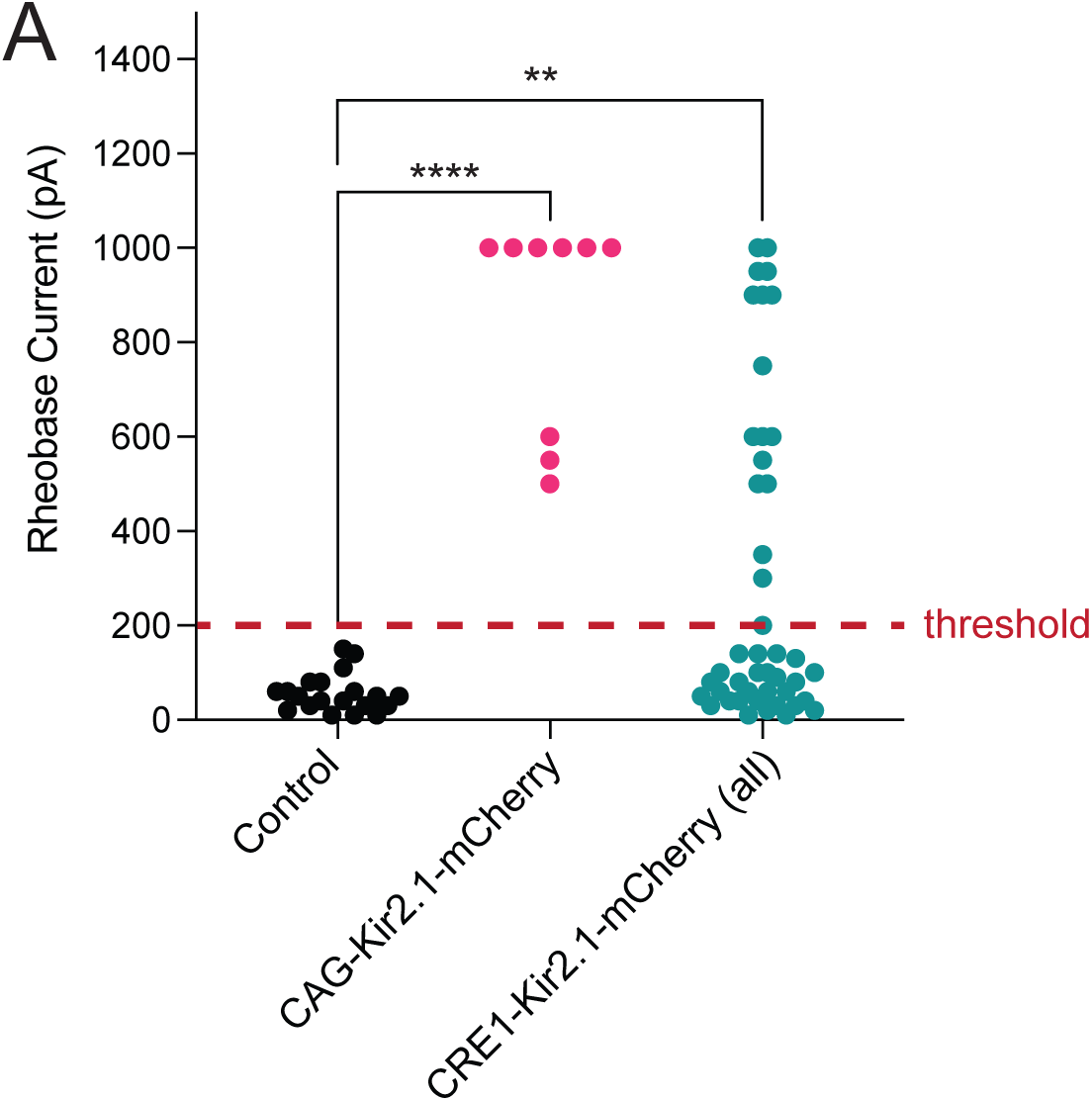
Functional applications of nociceptor-specific AAVs. **A:** Summary data showing the minimum current required to elicit an action potential (rheobase). Data is the same as in Figure 6C but also includes data from all CRE1-K_ir_2.1 neurons without functional stratification (>200 pA rheobase, red dashed line) to confirm K_ir_2.1 expression. Statistical significance was determined using a one-way ANOVA with multiple comparisons (* p < 0.05, ** p < 0.01, **** p < 0.0001). Sample sizes (Control, n = 20; CAG, n = 9; CRE1 total, n = 45) represent individual neurons recorded across at least 2 independent preparations.

## Acknowledgements

We would like to thank donors and donors’ families, without whom much of this study would not be possible. We also thank the NIH HEAL PRECISION Human Pain Network for its support and for fostering a highly collaborative research environment.

We thank Junning Case from the BWH Center for Cellular Profiling for assistance in experimental design and protocol optimization. We would also like to thank Rajesh Krishnan from The ARCND Flow Cytometry Core Facility for assistance in protocol optimization. We also thank the BWH Neurotechnology Studio. The authors thank Jesse Gillis from the University of Toronto for his advice on how to build PAIN-net. Schematics in Figures 1, 3, 4, and 6 were adapted using icons provided by <u>BioRender.com</u>. Finally, the authors would like to thank members of the Renthal Lab for their advice throughout the process and Hannah MacMillan for assistance with figure diagrams.

## Data and Code Availability

All custom code used to integrate data is available on Git (https://github.com/Renthal-Lab/painseq-multiome). Raw sequencing data and final objects will be deposited into GEO after manuscript acceptance.

## Author contributions

**L.S.H., P.B., S.A.B., and W.R.** designed, conceptualized, wrote, and reviewed the study. **J.W., L.J., M.X., L.Y., and L.S.H.** constructed AAVs. **P.B., S.A.B., and D.Z.** performed bioinformatics analyses. **E.S.** performed mouse and human DRG multi-omic profiling. **J.L., M.X., L.Y., E.G., C.A., L.S.H., K.P., E.K.W., and J.W.** performed and analyzed *in situ* hybridization. **L.S.H. and J.W.** performed single-cell sequencing. **J.L. and S.J.** performed iPSC experiments. **L.S.H., J.L., J.W., H.J.Y., and Q.L.** performed *in vivo* injections and procedures. **J.N.** and **B.W.** performed electrophysiology and associated analysis. **P.B., J.W., J.X.J.L., S.H. and W.R.** performed enhancer prioritization and library design. **D.G.** performed donor DNA sequencing and demultiplexing. **L.S.H., P.B., S.A.B., and W.R.** wrote the manuscript with input from all authors. **W.R.** supervised the research. All authors reviewed and approved the manuscript.

